# Microeukaryote metabolism across the western North Atlantic Ocean revealed through autonomous underwater profiling

**DOI:** 10.1101/2023.11.20.567900

**Authors:** Natalie R. Cohen, Arianna I. Krinos, Riss M. Kell, Rebecca J. Chmiel, Dawn M. Moran, Matthew R. McIlvin, Paloma Z. Lopez, Alexander Barth, Joshua Stone, Brianna A. Alanis, Eric W. Chan, John A. Breier, Michael V. Jakuba, Rod Johnson, Harriet Alexander, Mak A. Saito

## Abstract

Protists (microeukaryotes) are key contributors to marine carbon cycling, influencing the transfer of energy to higher trophic levels and the vertical movement of carbon to the ocean interior. Their physiology, ecology, and interactions with the chemical environment are still poorly understood in offshore ecosystems, and especially in the deep ocean. Using the Autonomous Underwater Vehicle (AUV) *Clio*, the microbial community along a 1,050 km transect in the western North Atlantic Ocean was surveyed at 10-200 m vertical depth increments to capture metabolic microeukaryote signatures spanning a gradient of oligotrophic, continental margin, and productive coastal ecosystems. Plankton biomass was collected along the surface of this transect and across depth features, and taxonomy and metabolic function were examined using a paired metatranscriptomic and metaproteomic approach. A shift in the microeukaryote community composition was observed from the euphotic zone through the mesopelagic and into the bathypelagic ocean. A diverse surface assemblage consisting of haptophytes, stramenopiles, dinoflagellates and ciliates was represented in both the transcript and protein fractions, with foraminifera, radiolaria, picozoa, and discoba proteins enriched at >200 m depth, and fungal proteins emerging in waters >3,000 m depth. In the broad microeukaryote community, nitrogen stress biomarkers were found in productive coastal sites, with phosphorus stress biomarkers in offshore waters where Saharan dust input is thought to supply iron and nitrogen. This multi-omics dataset broadens our understanding of how microeukaryotic taxa and their functional processes are structured along environmental gradients of temperature, light, macronutrients, and trace metals.

## Introduction

The western North Atlantic Ocean encompasses a diverse collection of ecosystems, including a subtropical gyre, a continental margin, and productive coastal waters, differing in temperature, nutrient availability, and organic carbon levels. The warm subtropical gyre waters are largely separated from cold, productive subpolar waters by the western boundary Gulf Stream current, with mixing occurring across the front ^1–3^ (Fig. 1). These distinct oceanographic properties select for the type of microorganisms present and their functional capabilities, with distance from shore and temperature acting as key influences shaping microbial communities ^4^. In oligotrophic gyre waters of the North Atlantic, nitrogen (N) limitation is a primary bottom up control on phytoplankton abundance, with phosphorus (P) and iron (Fe) additionally limiting or co-limiting growth ^5–7^. Little is understood about the metabolic status of microbial communities occupying transitionary waters between the subtropical gyre and the continental shelf, and how the differing nutrient supplies and physical dynamics influence phytoplankton physiology along this gradient.

**Fig. 1.**
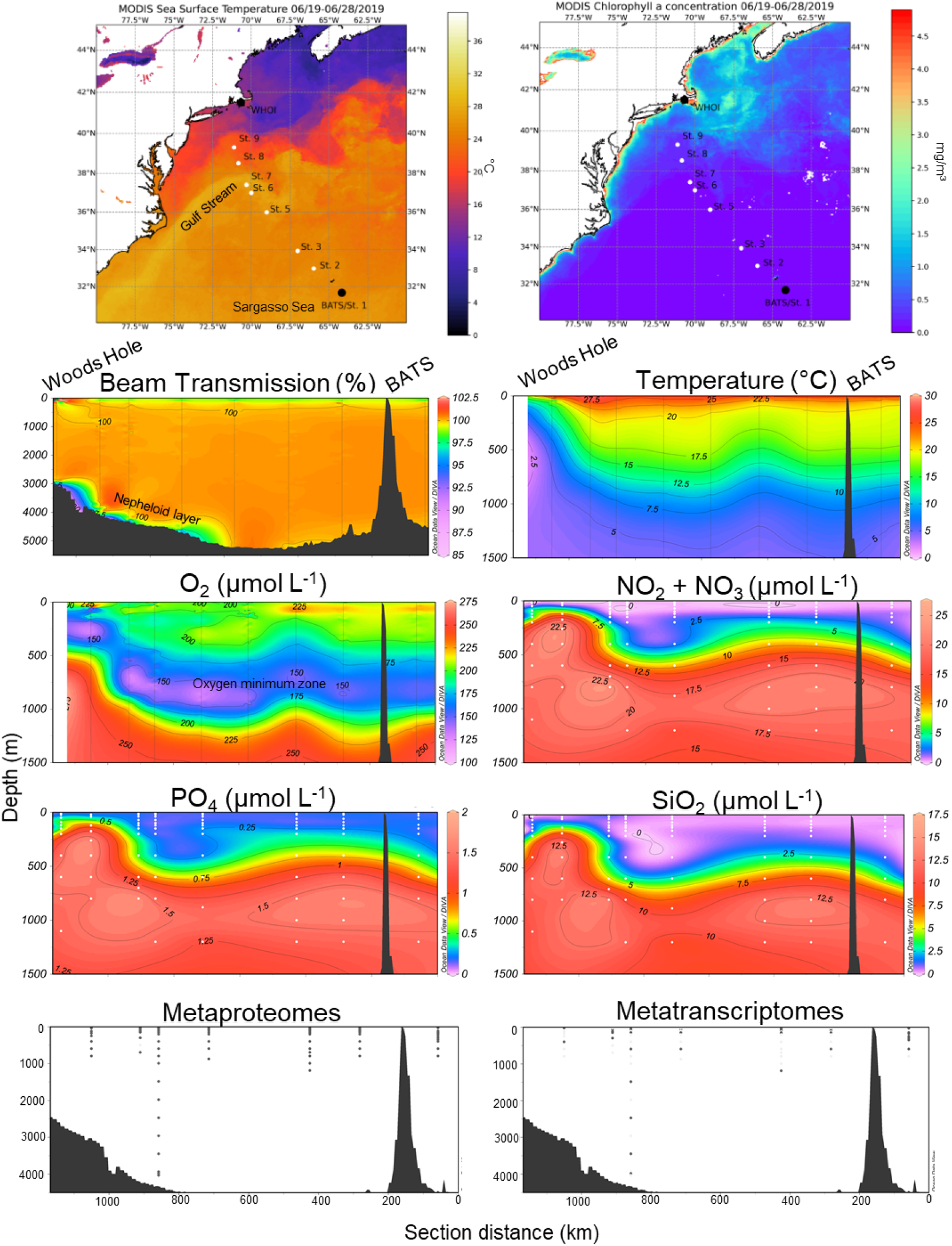
Transect sampling began at the oligotrophic site Bermuda Atlantic Time Series (BATS), and ended in coastal waters of the Northeast US continental shelf. near Woods Hole Oceanographic Institution (WHOI). Biological sampling was performed using the AUV *Clio* and additional parameters were collected via CTD rosette at each station. The surface transect is shown overlaying sea surface temperature and satellite-derived chlorophyll *a* concentrations between June 19-28th 2019, obtained from NASA MODIS. Continuous beam transmission (turbidity), temperature, and dissolved oxygen measurements were obtained via the ship CTD, and dissolved nutrients were measured from discrete samples from the top 1,500 m of the water column (see Supplemental Fig. 1 for full depth temperature and oxygen sections). Seawater samples collected by *Clio* for metaproteomes and metatranscriptomes were obtained from the depths shown in the bottom section plots.

The planktonic communities in the western North Atlantic experience strong seasonal forcings, subject to interannual and decadal-scale climate variability ^8^. A major source of nutrients to the North Atlantic is seasonally variable aerosol dust from the Saharan Desert ^9–12^, with most of the deposition in Bermuda occurring in the late summer ^13^. These deposition events introduce dissolved Fe and N to surface waters, which is rapidly consumed to support phytoplankton growth and may drive the system towards phosphorus (P) limitation ^14^. Toward the subpolar-influenced northeast US continental margin, overall phytoplankton biomass increases and the shelf/slope becomes a dominant source of Fe ^15^ ^16,17^. The offshore (subtropical) to coastal (subpolar) section thus represents a natural biogeochemical nutrient gradient (Fig. 1). The North Atlantic continental shelf ecosystem is also experiencing rapid warming, and the chemical and ecological ramifications of this for phytoplankton communities are still unclear ^18^.

Microeukaryotes, or protists, serve critical roles in the marine environment by performing primary production, transferring organic material to higher trophic levels, moving carbon to deeper layers of the ocean, and facilitating symbiotic relationships ^19^. Microeukaryotes are capable of strict autotrophy, strict heterotrophy, or a combination of the two trophic strategies through mixotrophy ^20^. Three decades of rich biogeochemical data collected at the Bermuda Atlantic Time-series Study (BATS) site located in the Sargasso Sea has established a baseline for phytoplankton production and community structure in this region ^21^. Surface phytoplankton communities are generally dominated by small cells including cyanobacteria and haptophytes ^22^, with cyanobacteria abundant in the winter and picoeukaryotes/nanoeukaryotes abundant in the spring, coinciding with a shoaling of the mixed layer and the annual North Atlantic spring bloom ^23,24^. Deep sea microeukaryote communities in this region are distinct from surface communities, and include *Acantharea* and *Polycystinea* radiolaria and *Euglenozoa* discoba ^25,26^. Although considerable progress has been made in cataloging microeukaryote diversity and ecology in the oligotrophic North Atlantic, we lack an understanding of 1) how these oligotrophic communities differ from those in continental margin ecosystems, and 2) which environmental factors could be responsible for changes in community structure and physiology along such transition zones.

Obtaining an accurate understanding of microeukaryote metabolism and their relationships to nutrient availability over large spatial expanses of the open ocean requires specialized sampling approaches. In particular, vertical profiling autonomous underwater vehicles (AUVs) capable of *in situ* filtration are particularly useful for the collection of microbial biomass ^27,28^. *Clio* is a specially designed AUV that actively transits in the vertically in the water column to a maximum depth of 6,000 m, drifts in a Lagrangian manner, and is outfitted with sensors for measuring chlorophyll *a* fluorescence, salinity, pressure, temperature, dissolved oxygen and optical backscatter ^27^. *Clio* can collect biomass by holding position at <5 cm vertical resolution ^27^, and therefore enables an unparalleled opportunity to characterize microeukaryote metabolism using a highly efficient vertical sampling scheme.

Here we present microbial community structure and metabolic profiles obtained with *Clio* along a western North Atlantic Ocean offshore-coastal transect, with a focus on microeukaryotes. The ecosystems along this transect were characterized by physicochemical parameters, macronutrients, pigments, and dissolved trace metals. Physiology and ecological interactions were inferred through complementary data-dependent-acquisition (DDA) mass spectrometry metaproteomics and short-read metatranscriptomics. Our results demonstrate that the microeukaryote communities were primarily composed of stramenopiles, dinoflagellates, and ciliates, which were detected throughout the water column. We identify several incidents of concordance and discordance between the transcript and protein pools investigated, which together construct a view of the epipelagic, mesopelagic, and bathypelagic microbial communities and their functional roles. Along the environmental gradient, the nutrient physiology of major microeukaryotic groups shifted with evidence of nitrogen stress in coastal waters and phosphate stress in the oligotrophic gyre. We furthermore examined the vertical structure of microeukaryote metabolism through the water column, revealing functional processes coupled to depth zone and co-occurrences among members of different taxonomic groups.

## Results & Discussion

This study region encompasses several nutrient regimes: a subtropical/oligotrophic ecosystem, a transitional continental margin region impacted by the Gulf Stream, and coastal productive waters receiving subpolar influence. Prominent oceanographic features captured include a persistent low oxygen zone, a sedimentary nepheloid layer along the continental margin, and deep chlorophyll maxima (Fig 1 & Supplemental Fig. 1). This transect is similar to the Western leg of the GEOTRACES GA03 North Atlantic Zonal Transect (NAZT), also known as Line W ^29^.

### Taxonomic composition of RNA and proteins

Transcripts were assembled using sequences derived from poly-A-selected mRNA, which enriches for eukaryotic members of the community. Unexpectedly, 23% of the annotated open reading frames (ORFs) belonged to prokaryotes (Fig. 2C), indicating that bacterial mRNA was not completely excluded in the library preparation process. The metatranscriptome was therefore determined to be suitable for the detection of eukaryotes and as a qualitative assessment of prokaryotes across the transect (Fig. 2). Eukaryotes were not similarly enriched during protein processing, and the metaproteomes therefore more accurately reflect the natural proportion of prokaryotes and eukaryotes in the water column (Fig. 2B). A notable feature revealed by the metaproteomes is that of the prokaryotes detected, cyanobacteria consistently peaked in relative abundance at the deep chlorophyll maxima (DCMs) throughout the transect (Fig. 2B), despite the relatively small number of cyanobacteria proteins predicted from the metatranscriptome (Fig. 2C). This is consistent with previous observations of elevated *Prochlorococcus* abundance at the DCM in the Sargasso Sea, where this organism thrives under warm, low macronutrient and low light conditions ^30^. For the majority of this analysis, we focus our investigations into the microeukaryotic component of this diverse microbial community. Additional features of the pelagic prokaryotic community of the Sargasso Sea determined with *Clio* have been reported elsewhere ^27^ and will be reported in a future study.

**Fig. 2.**
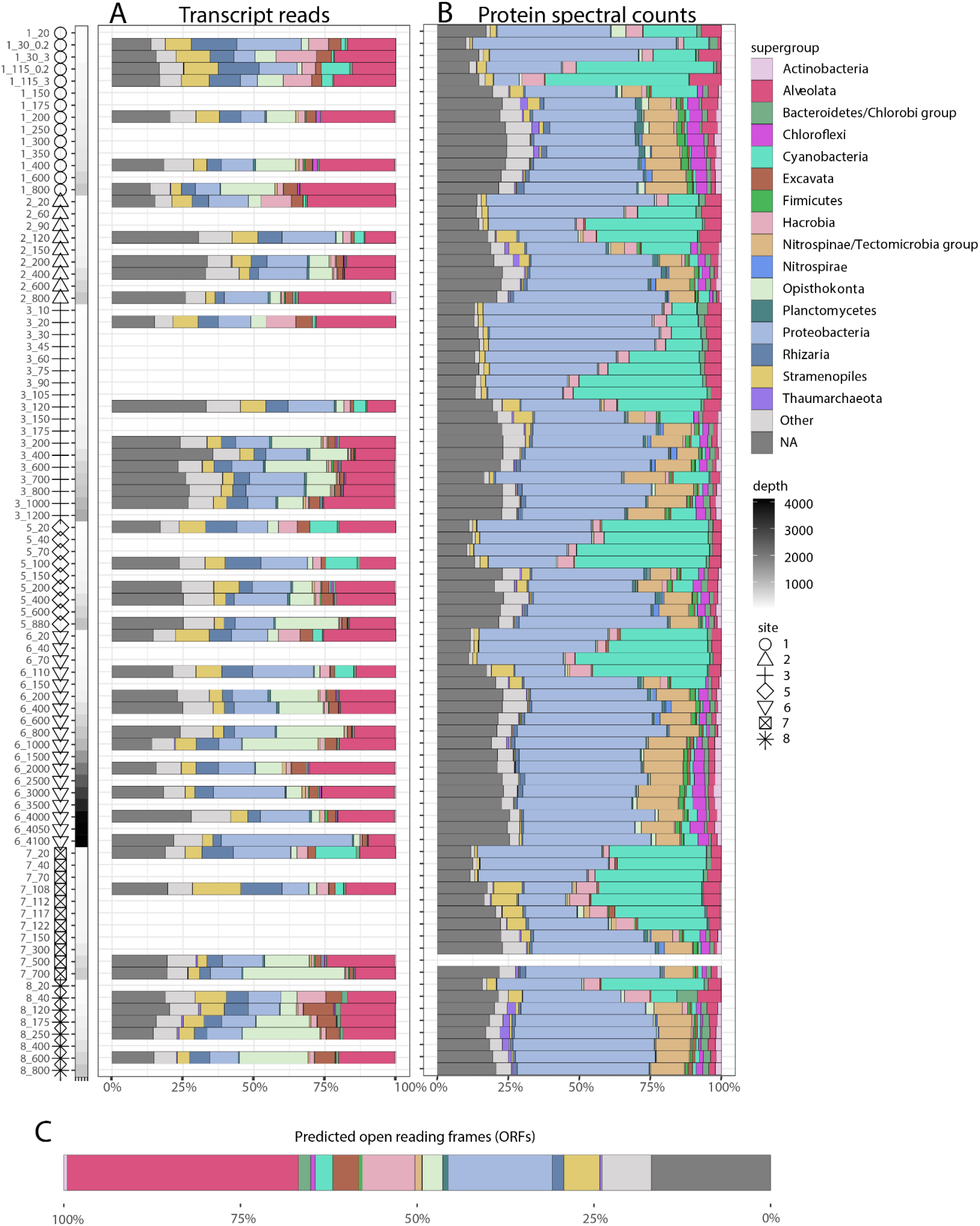
Relative abundance of transcript reads recruited to open reading frames (ORFs) (A), and exclusive proteomic spectral counts assigned to these ORFs (B) shown for the whole microbial community. All microbial groups with a EUKulele taxonomic annotation are depicted. Reads associated with unannotated ORFs are not shown. Each row represents a station and depth (“7_700” = St. 7, 700 m). Note that 30 m and 115 m surface samples at BATS (St. 1) were collected using McLane pumps instead of *Clio*, with separate 0.2-3 μm (“0.2”) and 3-51 μm (“3”) filter fractions. Both fractions are shown. NA represent microbes with lineage-conflicted taxonomic assignments unable to be resolved at this broad level. (C) The relative abundance of taxonomically annotated ORFs contained in the translated metatranscriptome database highlights the large percentage of unique alveolate (including dinoflagellate) and Proteobacterial ORFs, and the low number of cyanobacterial ORFs. Despite not having many ORF references, cyanobacteria were detected in relaticely high numbers in the metaproteomes in the deep chlorophyll maxima (B).

Of the total community detected on filters, approximately one quarter of the transcript reads and protein spectral counts were associated with predicted ORFs that could not be distinguished at the broadest classification level (“NA”; Fig. 2). These are ORFs that were similar to multiple, distantly related references in our taxonomic database such that they were not possible to confidently assign using our sequence homology and LCA classification approach ^31^.

The 10-200 m resolution sampling scheme of our metaproteomic approach allows for the visualization of strong vertical shifts in eukaryote community composition at both the supergroup level and at finer taxonomic levels. Surface communities were comprised of dinoflagellates (*Gymnodiniales*, unidentified marine alveolates [MALVs]), haptophytes (*Prymnesiales*), mixed stramenopiles (dictyophytes, diatoms, pelagophytes), ciliates (*Spirotrichea*) and chlorophytes (Fig. 3; Supplemental Fig. 2–5). The diversity of surface phytoplankton communities are supported by pigments measured from the water column demonstrating the presence of diatoms (fucoxanthin), haptophytes (19’Hex-Fucoxanthin and fucoxanthin), chlorophytes (chlorophyll *b*), and cyanobacteria (zeaxanthin and divinyl chlorophyll *a*) ^32^ (Supplemental Fig. 11). Dinoflagellates are also known to harbor many of the eukaryotic pigments, gained through secondary and tertiary endosymbiosis ^33^. Although pigments have long been used to decipher marine phytoplankton community composition, biomarker pigments may be variable within and among taxonomic groups, complicating efforts to directly relate them to taxonomic assemblages ^34^. The coherence among pigments, transcripts, and extracted protein pools are further discussed below (See *Quantitative estimates of biomass*).

**Fig. 3.**
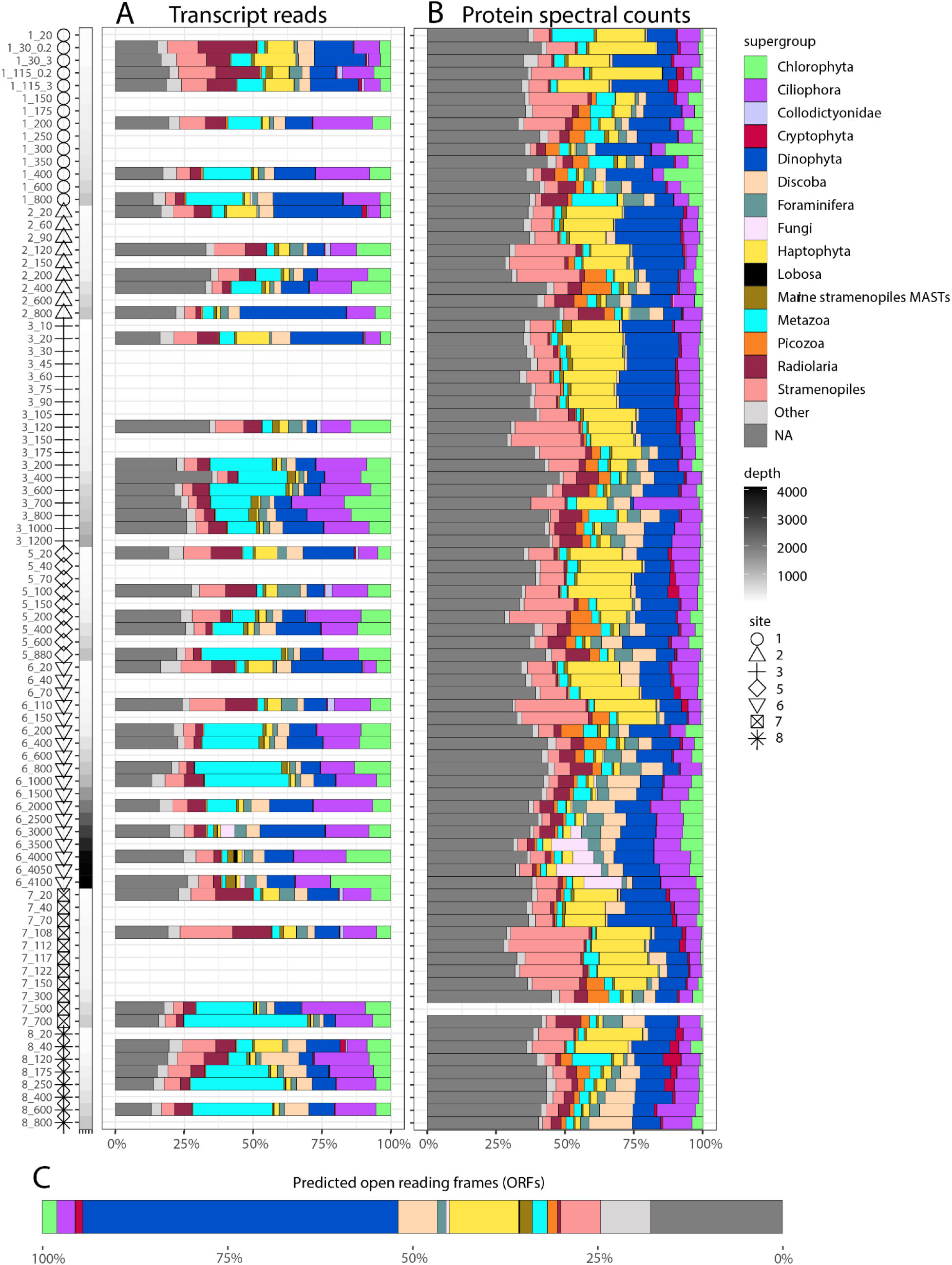
Relative abundance of transcript reads recruited to ORF predicted proteins (A), and exclusive proteomic spectral counts assigned to ORFs (B) subset within the eukaryotes. Each row represents a station and depth (“7_700” = St. 7, 700 m). Only microeukaryote transcripts having a EUKulele taxonomic annotation are shown. Reads associated with unannotated ORFs are not shown. Note that 30 m and 115 m surface samples at BATS (St. 1) were collected using McLane pumps instead of *Clio*, with separate 0.2-3 μm (“0.2”) and 3-51 μm (“3”) filter fractions. Both fractions are shown. NA represent microbes with lineage conflicted taxonomic assignments unable to be classified at the eukaryote supergroup level. (C) The relative abundance of taxonomically annotated eukaryotic ORFs contained in the translated metatranscriptome database is displayed as a single stacked barplot.

Mesopelagic communities >200m reflected a greater relative abundance of phaeophyte and pelagophyte stramenopiles, foraminifera (*Rotaliidae*), radiolaria (*Symphyacanthida),* discobans (*Euglenozoa*) and picozoa (Fig. 3, Supplemental Fig. 6–8). Many of these groups are similarly relatively abundant in the Sargasso Sea based on 18S rRNA V4 amplicons, with communities also shifting toward heterotrophic taxa below the DCM ^26,35^. Dinoflagellates, stramenopiles, fungi and radiolarians are furthermore present in the deep ocean sediment and are hypothesized to be important taxa involved in carbon export ^36,37^. These taxa contain autotrophic, heterotrophic, and mixotrophic members, and our deep omic signatures could reflect cells both passively sinking from the photic to aphotic zone, and resident communities adapted to the deep ocean. The vertical community shifts were more pronounced in the protein pool, whereas transcripts were more evenly distributed throughout the water column (Fig. 3A-B).

Although the proteomic assessment demonstrated heterotrophic microeukaryotes as key taxa in deep waters, foraminifera in the mesozooplankton size fraction (>500 μm) was in highest absolute concentration in surface waters along the continental margin, as determined via Underwater Vision Profiler (UVP) imagery (Supplemental Fig. 12). UVP imagery also supports radiolarian (*Collodaria, Phaeodaria,* and *Acantharea*) presence across the transect with highest concentrations in surface waters towards the coastline where plankton biomass accumulates (Supplemental Fig. 12). These cell counts differ from our protein profiles (Fig. 3B, Supplemental Fig. 12) and previously reported 18S rDNA composition ^26^, in which rhizaria of the Sargasso Sea notably increase with depth. These differences are likely due to rhizaria being dominant microeukaryotes in the mesopelagic where overall plankton biomass is low. It is important to note that the omics method captured microeukaryotes in the 0.2-51 μm size range while UVP captured mesozooplankton >500 μm, thus these methods are not directly comparable. Deep sea radiolaria in particular may additionally be smaller than commonly observed surface taxa or have sufficiently different morphology, complicating direct comparisons between imaging and omic data sets ^38^. In addition, the radiolarian transcripts and proteins detected could be derived from the <5 μm gamete-like swarmer reproductive stage ^39^ captured on *Clio* filters, while images are biased towards the larger adult stage. Radiolaria are hypothesized to sink down below the mesopelagic zone to complete their reproduction cycle ^39^, which is consistent with radiolarian proteins observed >200 m along the transect (Fig. 3B). It is also worth noting that we are less confident in fine-scale rhizaria taxonomic assignments given their poor representation in existing databases ^31^.

Microeukaryote community composition continued to shift through the mesopelagic and into the bathypelagic. *Ascomycota* fungal proteins were enriched below 3,000 m at St. 6, the only site where bathypelagic water was collected (Fig. 3B). Fungi are important microeukaryotes driving biogeochemical cycling and are poorly understood in marine systems ^40^. These results are consistent with 18S rDNA data that show that this group comprises important members of the global bathypelagic zone, along with rhizaria (foraminifera and radiolarians), stramenopiles, and alveolates (including dinoflagellates and ciliates) ^41^. These fungal proteins may represent small cells or structures (< 51 μm), similar to small fungi detected in the 3-10 μm filter fraction in coastal ecosystems ^42^. Large fungal structures (>51 μm) would have been missed in this omic analysis unless captured as cell debris during remineralization or broken apart during filtration through the 51 μm membrane. Approximately half of these annotated fungal proteins above the seafloor (4,050-4,100 m) were aldehyde dehydrogenases, involved in a number of cellular processes including detoxification and biosynthesis pathways ^43^. A benthic nepheloid layer was observed at this location as determined with beam transmission (turbidity), appearing at ~4,000 m (Supplemental Fig. 13). A nepheloid layer in this region has been previously characterized on the GEOTRACES NAZT expedition along the Line W transect, establishing elevated suspended particulate matter above the continental margin, largely composed of lithogenic material with slight enrichment in particulate organics ^44^. Particulate trace metals concentrations are additionally elevated in this nepheloid layer, potentially serving as a metal source to bathypelagic microbes ^45,46^. Fungi are consumers of organic detrital material ^47,48^, are hypothesized to contribute to methane and nitrogen cycling ^49^, and are active degraders of carbohydrates and proteins throughout the water column and in the subseafloor ^50–52^. It is unclear whether fungi benefit from enhanced particulate material in the nepheloid layer above the seafloor, or are generally abundant at these depths of the western North Atlantic. Additional deep ocean surveys are required to further understand the vertical spatial extent of pelagic fungi in the bathypelagic and their relationship to nepheloid layers ^40,53^.

### Concordance between transcript and protein pools

There was no linearly agreement between individual transcript and proteins on the ORF level (r^2^=0.005), but there was strong agreement when aggregated to the Kyoto Encyclopedia of Genes and Genomes (KEGG) Ortholog functional level (r^2^=0.86) (Supplemental Fig. 14). We interpret this to indicate certain ORFs receive peptide spectral counts that transcripts do not, and vice versa, potentially due to redundant taxonomic features and the nature of assigning peptide spectrum matches to representative protein groups. But, when aggregated to higher functional classification, processes that were relatively abundant were reflected in both pools.

We next investigated dataset-wide relationships between individual microeukaryote transcripts and proteins. To achieve this, a partial least squares (PLS) regression was performed on genes of interest (Fig. 4). Nitrogen biomarkers showed linear agreement, with a nitrate transporter (NRT) protein positively correlated with NRT, nitrate reductase, ferredoxin-nitrite reductase and glutamate synthase transcripts. These transcripts were also positively correlated with cytochrome b_6_f complex proteins, consistent with nitrogen utilization and photosynthesis co-occurring in the upper water column. Transcripts and proteins of the respiratory protein NADH dehydrogenase furthermore positively correlated, specifically subunits 2 and 3. Interestingly, ferric chelate reductase transcripts, which reduce Fe^3+^ to Fe^2+^ for cellular transport, were also positively correlated with NADH dehydrogenase subunit 2, which may be involved in Fe transport via the ferric reductase complex in surface waters ^54^. NADH dehydrogenase proteins demonstrated some of the strongest positive relationships with transcripts also expressed in surface waters, including a zinc transporter (ZIP2), flavodoxin, carbonic anhydrases, and the inorganic phosphate transporter PHO84 (Fig. 4). Negative correlative relationships were also identified, driven by water column depth where metabolic processes were occurring. For example NRT proteins were detected in surface waters where cells were investing in nitrogen uptake to fuel photosynthetic growth, while ferredoxin and Cu/Zn SOD transcripts in deep water where they presumably supported metal-dependent heterotrophic metabolism. Conversely, surface transcripts related to nitrogen assimilation and photosynthesis (flavodoxin, nitrate reductase) were negatively correlated with peptide/nickel transport system proteins that were primarily detected in the deep ocean. In addition, many transcript/protein pairings showed no strong linear relationships. These include ferritin, Cu/Zn superoxide dismutase, and the inorganic phosphate transporter (Fig. 4). These transcripts and proteins could have large inventories with more stable expression compared to dynamically regulated processes such as NRT ^55^.

**Fig. 4.**
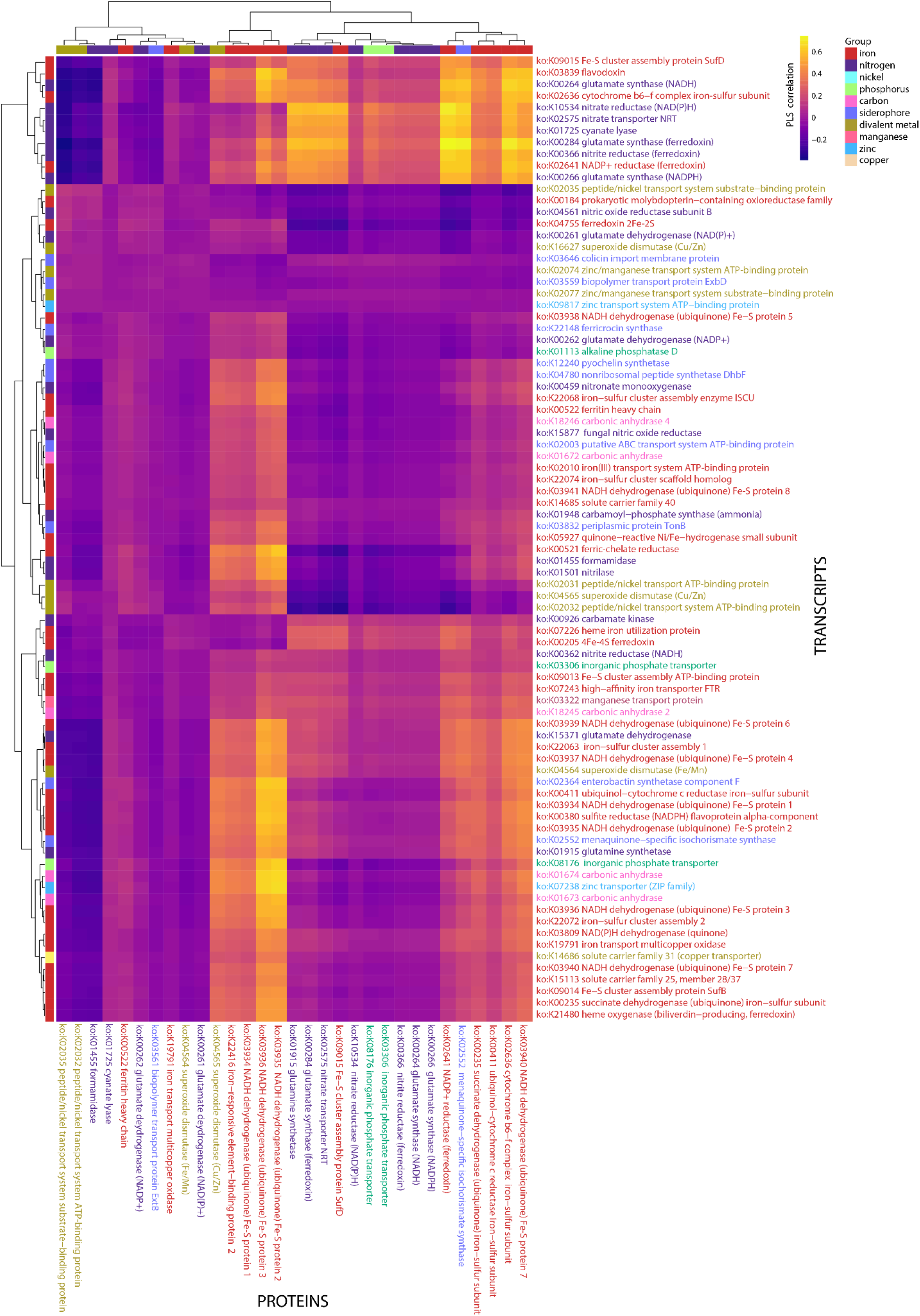
Partial least squares regression analysis showing the relationship between nutrient and metal-related transcripts and proteins. Transcripts are labeled as rows (horizontal), proteins as columns (vertical), and clustered based on similarity. Annotations are color coded by functional categories (legend “Group”). Note: more KEGG Orthologs were detected in the metatranscriptome than in the metaproteome, resulting in more rows on the x-axis.

In terms of taxonomic agreement, in several cases, transcripts and proteins differed in the taxa captured and resulting community distributions. In surface waters, haptophyte proteins were well-represented, reflecting 17% of eukaryotic proteins <200 m (Fig. 3B), but transcripts suggest they were minor members of the surface community, only comprising 7% of eukaryotic reads (Fig. 3A). Similarly, metazoan transcripts were relatively abundant (Fig. 3A), with most transcript reads belonging to a single sponge reference in the EukProt database (*Vazella pourtalesii;* Supplemental Fig. 10), while metazoa were comparatively minor constituents of the protein pool (Fig. 3B). Differences were additionally apparent at finer taxonomic levels. Silica skeleton-containing *Polycystinea* radiolarian transcripts were detected throughout the water column, yet proteins were mainly detected in the mesopelagic and largely represented by strontium sulfate-containing *Symphyacanthida* instead ^56^ (Supplemental Fig. 8). Transcripts indicated a diverse ciliate population including *Armophorea, Oligohymenophorea*, and *Spirotrichea*, while proteins indicated that the ciliate population was nearly exclusively composed of *Spirotrichea* (Supplemental. Fig. 5). On the metaproteomic side, these patterns could in part be due to shared peptides between closely related taxa masking diversity ^57^, and/or not detecting rare peptides that are sequenced in transcriptomes. In the transcript fraction, certain organisms may be relying on post-transcriptional modification for protein regulation, resulting in a larger transcript pool (Fig. 3A).

In addition to these taxonomic distinctions between biochemical pools, we observed an unexpected difference in the proportion of the eukaryotic communities able to be confidently classified ^31^. Approximately 21% of transcripts on average were associated with ORFs unclassified at the supergroup level, while an average of 38% protein spectral counts were associated with lineage-conflicted ORFs (“Not Annotated (NA)”; Fig. 3). These lineage-conflicted proteins were composed of a higher relative abundance of cytoskeletal proteins (Supplemental. Fig. 15). We attribute the large portion of lineage-conflicted eukaryotic ORF proteins to the collection of evolutionary conserved protein machinery with high sequence similarity across supergroups, such as exoskeleton components ^58^, potentially with large inventories and slow degradation rates. These proteins may belong to a collection of genomically diverse eukaryotes sharing conserved proteins, or could instead belong to one or a few taxa dominant in the environment that are not included in the taxonomic database used here. It also is possible *Clio* captured these abundant and conserved proteins shed from organisms across trophic levels in the ecosystem as “eProtein”, similar to “eDNA” ^59^, in addition to collecting biomolecules from intact cells. It is unlikely that the DDA mass spectrometry method was the reason behind this bias in conserved proteins. Although taxonomically conserved peptides may be abundant in extracted protein pools, a dynamic exclusion approach was used, which limits continued fragmentation of a given parent ion.

Instances of protein and transcript concordance are insightful, as transcripts and proteins are not expected to uniformly agree given that the two pools have inherently different cellular concentrations, degradation times, and regulatory controls. Proteins can be 30,000 times more abundant in cells than transcripts ^60^ and reflect cell activity, whereas transcripts are sensitive to rapid changes in the environment and subject to post-transcriptional regulation processes that may prevent transcripts from representing cell phenotype. Importantly, there are key differences in the half lives of these molecules. Transcripts are quickly produced and degraded in cells, and decrease once protein needs are met, whereas proteins are generally more stable though have highly variable turnover times ^61^. Hence, proteins and transcripts are distinct molecular pools that are not universally expected to correlate. In environmental data sets, the comparison between these two pools is further complicated by the differences in functional and taxonomic coverage (Fig. 3A-B), potentially due to filtration time, extraction, amplification, or analytical biases. Nevertheless, environmentally sensitive transporters and enzymes that accumulate in both transcript and protein fractions may indicate a cellular effort to rapidly maximize substrate uptake, and results in a strong signal detected in both molecular fractions, as predicted in a recent modeling study using mRNA and protein turnover times ^55,62^.

Other possible explanations for the observed discrepancies in our dataset between molecular pools include lineage-specific loss of proteins or RNA during the extraction, amplification, or analytical steps ^63,64^, and/or a biological distinction in their biochemical inventories with differences in RNA/protein ratios among taxonomic groups ^65^. It is furthermore probable that as biomass accumulates on filters during *in situ* filtration, a portion of cells lyse under pressure ^66^. This may lead to a loss in soluble RNA and proteins, but the retention of membrane-bound proteins along with particulate material on filters, thus contributing to observed differences between protein/RNA fractions. Large, fragile eukaryotes with gelatinous structures may be especially vulnerable to cell rupture during filtration. Both protein- and RNA-based community composition assessments may also be biased by inefficient zooplankton feeding and host-specific viral lysis, leading to the leakage of organic material from certain microeukaryotic groups ^67^. Both fractions may also be influenced by growth rate, with faster growing eukaryotic cells producing more proteins and especially more RNA ^68,69^, thereby reflecting higher relative abundance in the community. A multitude of analytical and biological factors therefore contribute to the taxonomic profiles observed here, which may differ from assessments based on bulk seawater cell densities. These results highlight that a combination of taxonomic metrics is more insightful than any one individually in reconstructing microbial community composition.

### Quantitative estimates of biomass

Biomass distributions were assessed using three metrics: semi-quantitative eukaryotic transcript estimates (copies L^−1^), total extracted protein (μg L^−1^), and total summed pigment concentrations (ng kg^−1^) (Supplemental Fig. 16–17). The highest levels of transcripts were generally measured in surface waters (St. 2-6), with St. 3 in particular having the highest detected concentrations of transcripts (Supplemental Fig. 16). Just prior to our occupation, this site had pronounced doming of the thermocline indicative of a cyclonic eddy, which causes upwelling of nutrient rich deep water into the surface layer ^70^. In contrast to the transcripts, pigments consistently showed highest concentrations at the deep chlorophyll maxima (Supplemental Fig. 11), which may reflect both increases in biomass and increases in chlorophyll *a* content per cell under low light conditions ^71^. As such, surface transcripts and pigments were not linearly correlated (r^2^ = 0.05) (Supplemental Fig. 17). Transcript copies and extracted protein however generally showed stronger agreement along the transect (r^2^ = 0.44), with both pools enriched in surface waters and decreasing in the aphotic zone (Supplemental Fig. 17). Extracted protein likewise linearly varied with pigment concentrations (r^2^ = 0.40), demonstrating that protein pools can be useful estimates of plankton biomass (Supplemental Fig. 17). Observed disconnects among these measurements may be partially explained by protein and pigment pools containing prokaryotes (Fig. 2, Supplemental Fig. 11), whereas the transcript data selected against prokaryotes by use of poly-A enrichment. In addition, growth dynamics could influence RNA levels, with transcripts potentially more concentrated in surface waters where cell growth is faster, for example following eddy-induced upwelling at St. 3, but with microzooplankton grazing likely preventing phytoplankton biomass from accumulating.

At the St. 8 coastal site, pigments, total extracted proteins, and eukaryotic transcripts all peaked in surface waters (40 m), where the thermocline shoaled and CTD chlorophyll fluorescence spiked (Supplemental Fig. 1), suggesting elevated biomass. A previous survey along a similar Sargasso Sea-coastal transect also showed highest chlorophyll *a* biomass north of the Gulf Stream where temperatures decreased, in which large plankton (>10 μm) were elevated in abundance ^30^. In particular, we observed the highest levels of fucoxanthin pigments at this site. There was a disconnect with omics data, with the relative abundance of stramenopile/diatom transcripts and proteins not similarly increasing at this location compared to other nearby sites (Fig. 3, Supplemental Fig. 5). This could suggest a limited ability of omics-based community composition data to reflect bulk changes in biomass.

Metatranscriptomics has been demonstrated to generally recapitulate phytoplankton groups using microscopy-based observations ^72,73^, but both molecular pools and pigment concentrations may be influenced by genetic factors and/or cell physiology, and therefore deviate from microscopy or image-based cell concentrations. The use of several RNA standards may produce more reliable quantitative transcript measures. Additional direct measurements of biomass (e.g. flow cytometry, microscopy) are recommended in future studies to further establish quantitative links between omics and cell densities, although these methods each have their own strengths and weaknesses. For example, flow cytometry excludes >50 μm eukaryotes, and traditional microscopy misses small picoplankton. Additionally, the size of organisms needs to be taken into account, as large organisms reach lower absolute cell concentrations given their larger volume, but still contribute to the biomolecule pool, perhaps even disproportionately. These results demonstrate that there is no perfect method for capturing eukaryotic community biomass, and multiple approaches are required to reconstruct community dynamics. Recommended methods will vary depending on the question of interest and the community members of interest.

### Eukaryotic metabolism between depth zones

Nonmetric multidimensional scaling (NMDS) was performed on taxonomically and functionally annotated transcripts and proteins to examine the spatial structure of microeukaryotic communities. Depth overshadowed lateral effects in both molecular fractions (Supplemental Fig. 18), with surface (< 100 m), upper water column (100-400 m), and deepest communities (400-4,100 m) separating in ordination space.

Co-occurrence networks highlighted taxonomic associations and linked depth-dependent functional processes. The transcript network consisted of 21,347 nodes, or eukaryotic supergroups with KEGG functional annotations, and 21,346 edges, or interactions between nodes. These nodes clustered into 156 modules (Supplemental Fig. 19). The protein network contained a smaller set of annotated ORFs, and consisted of 2,938 nodes and 2,937 edges which clustered into 56 modules (Fig. 5). Whereas many modules were taxonomically diverse, some were dominated by one or few supergroups (Fig. 5A). The large number of modules represent distinct sets of linked metabolic processes, and may represent different taxonomic groups at a fine taxonomic level, or co-regulated functional processes. These co-occurrences may be a result of similar niche space occupation, endosymbioses, predator-prey dynamics, or animal microbiome communities ^74,75^.

**Fig. 5.**
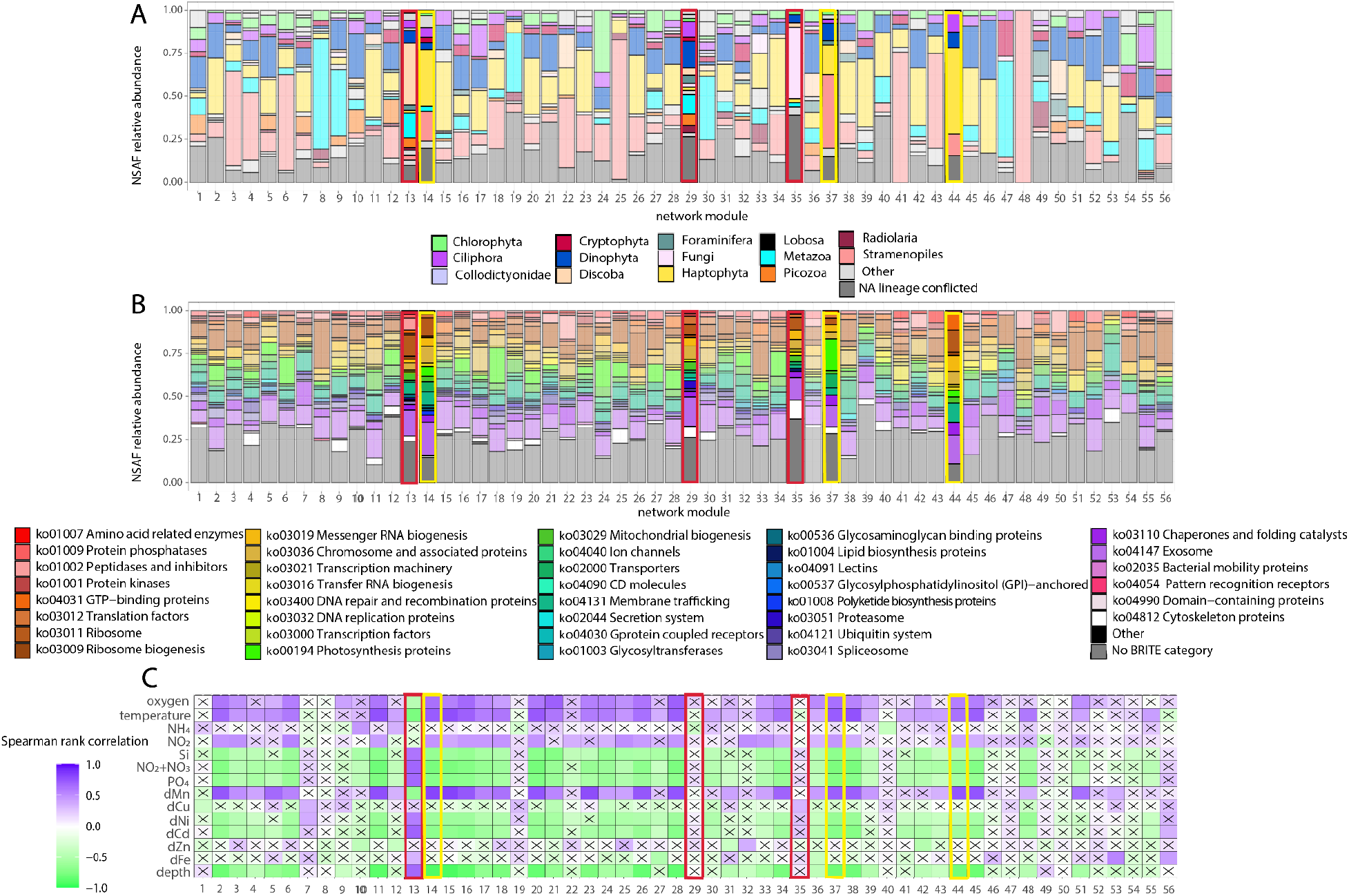
Taxonomic (A) and functional (B) composition of protein network modules. NSAF counts were summed to the supergroup and KEGG annotation level (e.g., Chlorophyta_ko:K00053). KEGG IDs were grouped into BRITE categories for a broader functional classification. KEGG IDs not included in BRITE categories are shown in dark gray. (**C**) Environmental context for the modules was determined by calculating eigengene values and correlating module eigengene values with environmental data. Insignificant Spearman rank correlations are denoted with an “x” (*p* > 0.05). Examples of surface ocean modules are highlighted in yellow, and deep water modules are shown in red, and are opaque for visibility in A&B.

The network modules showed strong representation of haptophytes, stramenopiles, and dinoflagellates, with several modules dominated by the heterotrophic taxa discoba, fungi, or metazoa. In contrast to other groups, metazoa were not relatively abundant in the protein pool (Fig. 3A-C), yet still comprised a large portion of certain modules, potentially reflecting consistent interactions with protists (Fig. 5A). The number of repeated KEGG Orthology genes (derived from different supergroups) contained in modules was low, ranging from 1-4, indicating that modules did not group by individual KEGG function. However, at a coarser functional level (KEGG BRITE category), differences in metabolic composition were observed. Several modules with an enrichment in photosynthesis were associated with higher relative abundances of photosynthetic taxa such as stramenopiles and haptophytes (Fig. 5B). These modules were positively correlated with oxygen, temperature and dMn, indicating elevated abundance in surface waters (Fig. 5C, Supplemental Fig. 19C).

Eight modules in the transcript network (Supplemental Fig. 19) and one in the protein network (Fig. 5) were significantly negatively correlated with temperature, and reflect deep ocean metabolism. These modules, along with several others showing negative (non-significant) correlations with surface parameters, were enriched in heterotrophic taxa such as discoba and foraminifera, and reduced in photosynthetic functionality. In particular, the deep ocean protein module 13 was mostly composed of discoba, metazoa, dinoflagellates, and picozoa, and contained a number of peptidases involved in amino acid catabolism (Fig. 5). Other protein modules were significantly positively correlated with temperature, yet negatively correlated with total dissolved Fe (dFe) (modules 27, 36 [Fig. 5]), and proteins in these modules included chlorophyll components (ascribed to chlorophytes, prasinodermophytes), and nitrate transporters (haptophytes). These modules therefore could reflect surface communities engaging in photosynthesis and assimilating macronutrients, thereby drawing down dFe concentrations to facilitate growth.

### Nitrogen, phosphorus and iron metabolism along the lateral gradient

Transcripts and protein biomarkers of carbon fixation, nutrient status, and metal metabolism were surveyed across the offshore-coastal transect and aggregated (summed) across all eukaryotic groups to investigate bulk metabolic fingerprints. They were evaluated in terms of absolute concentrations, either transcript copies L^−1^ or protein spectral counts L^−1^. Rubisco concentrations were higher closer to the coastline, and belonged to dinoflagellates, haptophytes, stramenopiles, cryptophytes, and unclassifiable eukaryotes (Fig. 6, Supplemental Tables 7-8). However, they peaked at different coastal locations, with Rubisco transcripts elevated in subsurface Gulf Stream waters (St. 7) with 12,000 transcript copies L^−1^, whereas Rubisco proteins reached a maximum of approximately 850 protein spectral counts L^−1^ in coastal surface waters of St. 8, which aligns with maximum pigment concentrations at St. 8 (Supplemental Fig. 11). This could represent differences in transcript/protein cellular ratios or turnover times. Growth rates of the phytoplankton populations between these two sites may also play a role, with higher RNA/pigment ratios in surface waters of the Gulf Stream supporting the possibility of elevated growth rates compared to coastal waters (Supplemental Fig. 11 & 16), resulting in higher Rubisco transcript abundance at St. 7.

**Fig. 6.**
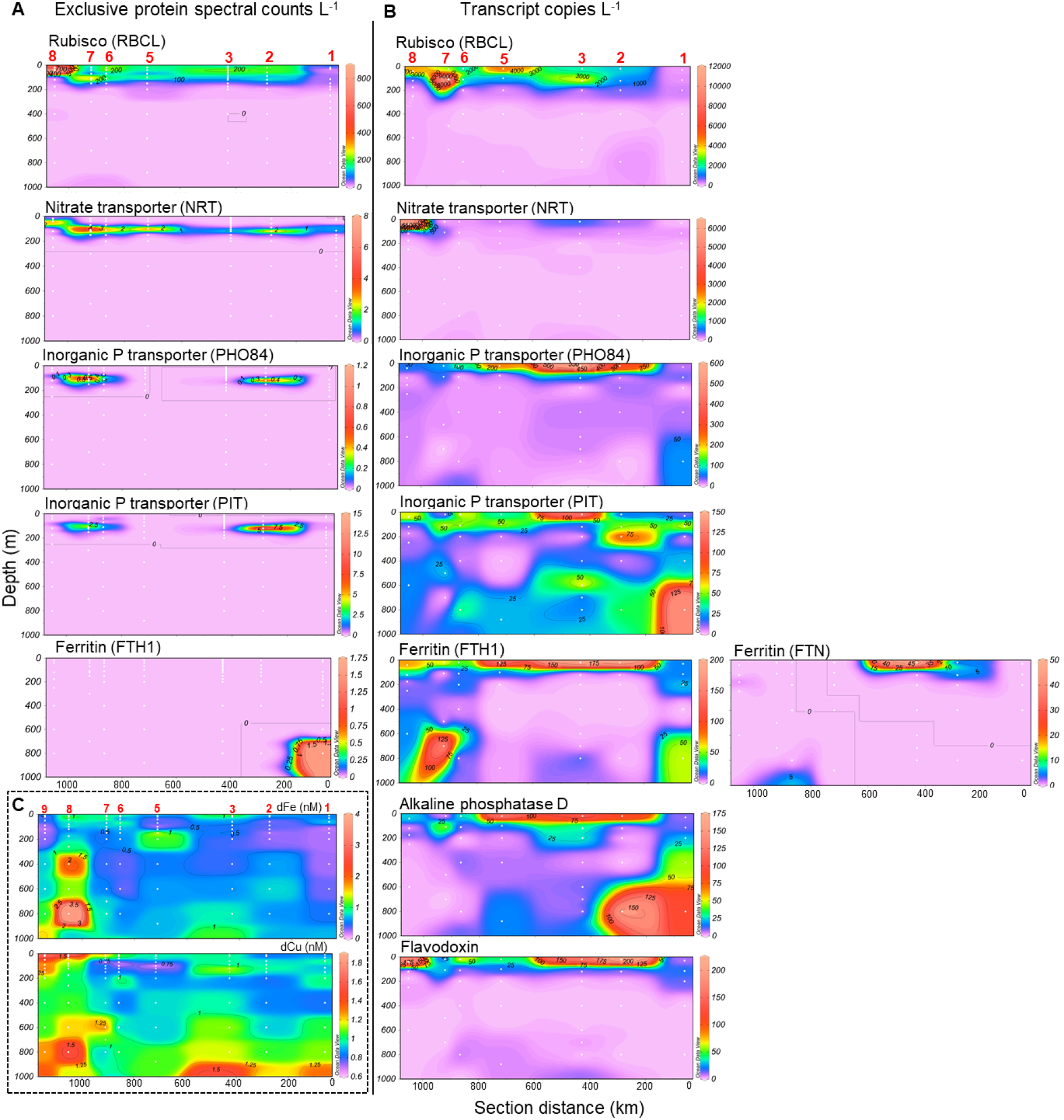
Protein spectral counts L^−1^ (A) and transcript copies L^−1^ (B) for functional processes of interest in the upper 1,000 m. Eukaryotic ORFs were summed to the supergroup level to examine total transcript read counts and protein spectral counts associated with a given KEGG functional gene. Two different KEGG-annotated ferritin proteins were detected in the transcripts (FTN = K02217, FTH1 = K00522). Eukaryotic alkaline phosphatase D and flavodoxin were only detected in the transcript fraction. (See Supplemental Tables 7-8 for individual open reading frame annotations). (C) Dissolved Fe and Cu concentrations in the top 1,000 m of the water column. Station numbers are labeled in red. Note that biomass for omics was not collected from the coastal site St. 9.

Macronutrient stress biomarkers suggest N stress occurred towards the continental shelf while P stress was apparent in offshore, oligotrophic gyre waters (Fig. 6). The nitrate stress biomarker NRT is a nitrate transporter in phytoplankton, upregulated under nitrogen starvation ^76^. NRT proteins reached highest levels in subsurface waters of the Gulf Stream (St. 7), and NRT transcripts peaked at the surface at the most coastal site (St. 8), suggesting that nitrogen may have been limiting phytoplankton growth in these coastal communities. NRT also belonged to dinoflagellates, haptophytes, stramenopiles, cryptophytes, and unclassifiable eukaryotes (Supplemental Tables 7-8).

Phosphate transporters showed a different distribution, with inorganic phosphate transporter PHO84, a major facilitator superfamily membrane transporter part of the Phosphate H^+^ Symporter family activated under phosphate stress ^77^, having highest transcript concentration at the surface along the middle of the surface transect (Fig. 6). These belonged to dinoflagellates, haptophytes, stramenopiles, discoba, centroplasthelida and unclassifiable eukaryotes. A second inorganic phosphate transporter (PiT) was detected as a minor signal in surface transitionary waters, but with transcripts most highly expressed in deep waters offshore, reflecting heterotrophic metabolism or sinking euphotic zone cells. PiT belonged to dinoflagellates, haptophytes, stramenopiles, cryptophytes and ciliates. Protein concentrations for both the PiT and PHO84 phosphorus transporters reached maximum concentrations in subsurface oligotrophic waters (St. 2) and in the Gulf Stream (St. 7), but were undetected at other sites. Alkaline phosphatase D, which is a metalloprotein that converts organic phosphorus to inorganic phosphorus, was only detected in the transcript fraction and was highly expressed in deep water at St. 1 & 2 (800 m), with a surface signal apparent in transitionary waters, matching PHO84 gene expression pattern. These transcripts were found in phototrophic taxa (dinoflagellates, stramenopiles, haptophytes) and heterotrophic taxa including choanoflagellates. Alkaline phosphatase has been demonstrated to be important in the subtropical Atlantic where inorganic phosphate concentrations are low, and enzyme activity increases with Saharan dust addition ^78^. Although not detected in eukaryotic proteins with the functional database used here, additional investigations into the prokaryotic fraction could provide insights into broader microbial use.

These omic signatures are consistent with a previous study conducted along a similar transect in the western North Atlantic in early spring, in which the inorganic N:P ratio of surface seawater was above Redfield ratio (16:1) in the oligotrophic gyre, and dropped closer to and below 16:1 north of the Gulf Stream, indicating the potential for phosphorus and nitrogen limitation in the oligotrophic gyre and coastal waters, respectively ^30^. However, there are strong seasonal shifts in N:P at BATS, with summer ratios dropping below 16 ^30^, and nitrate concentrations below detection limit for much of the year apart from deep mixing spring periods. Indeed in our summer analysis, nitrate+nitrite and phosphate concentrations were consistently low throughout the surface of the transect, with no clear transition in the N:P ratio towards the coastline (Fig. 1). As this survey was completed in early summer, it is possible that bioindications of nitrogen stress eventually appear in the oligotrophic gyre into the late summer and early fall, and a temporal analysis would be beneficial to further tease apart seasonal variations in nutrient physiology.

In addition to macronutrients, metalloproteins were also evaluated with respect to dissolved trace metal concentrations. The iron storage protein ferritin functions to maintain iron homeostasis ^79,80^ and is commonly elevated in diatoms under Fe enrichment ^81,82^. Two different ferritin genes were identified: “ferritin heavy chain” (FTH1: K00522) and “ferritin” (FTN: K02217). Although the only FTH1 protein signal (belonging to cnidarian metazoa) was detected in deep water, both FTN and FTH1 transcripts (comprised of metazoa, ciliates, foraminifera, telonemia, dinoflagellates, chlorophytes, streptophytes and unclassifiable eukaryotes) were found both in the center of the transect and in deep water (Fig. 6A-B). Transcripts of flavodoxin, an Fe-independent photosynthetic electron acceptor typically diagnostic of Fe stress ^83^, were abundant in surface transitionary waters alongside ferritin transcripts. These flavodoxin transcripts belonged to dinoflagellates, haptophytes and cryptophytes. These spatial patterns of ferritin and flavodoxin coincide with elevated dFe and dCu in the center of the transect compared to at BATS, which could be sourced from aerosol dust (Fig. 6C). High Fe availability would presumably lead to Fe sufficiency in cells, and some phytoplankton continue to rely on flavodoxin even under Fe enrichment ^84^ as an evolved strategy in oligotrophic, low Fe ecosystems. The addition of aeolian-sourced Fe and N may promote P stress ^14^, which is consistent with high PHO84 and alkaline phosphatase D signals alongside high FTH/FTN, FLDA and low NRT signatures in transitionary surface waters. This spatial pattern supports P being an important driver of microeukaryotic growth and community diversity in the Sargasso Sea ^35,85,86^.

### Future Directions

The data presented in this study leveraged sampling efficiencies enabled by coupled use of an autonomous underwater vehicle with concurrent shipboard sampling. Using AUV *Clio*, an average of 2,320 L of seawater was filtered per dive, concentrating biomass at 8-18 depths per station along a 1,050 km transect in 11 days, providing mid- and full-depth metabolic profiles (Supplemental Table 6). The average dive duration was 12.7 hours and sampling range was 10-4,100 m; during this time, other deck operations using the ship’s J-frame could continue, including trace metal rosette, CTD, and complementary *in situ* McLane pump deployments. Typical *in situ* underwater pumps are attached to the ship’s winch line and pump water at specific depths, needing manual adjustments for subsequent biomass collection at additional depths, and requiring the ship to wait on station until filtration is complete. Autonomous collection instruments such as *Clio* are thus highly efficient for oceanographic surveys requiring large filtration volumes. As station time is a major constraining factor in the design of sectional sampling expeditions, the efficiencies demonstrated in this study have implications for future study design. *Clio* is particularly valuable in efforts to jointly sample chemical parameters and omics material, as is the vision of the international *BioGeoSCAPES* program (https://biogeoscapes.org/), and can provide insights into microbial metabolism within physical and chemical features, such as nepheloid layers, oxygen minimum zones, and transitions in water masses. Future expeditions will benefit from further integrating *Clio* sensor data, and testing bulk seawater collection by *Clio* for trace metal quantification. Such advances may obviate simultaneous CTD deployments, and *Clio* alone could be used for multi-parameter collection.

## Conclusion

Here we present microeukaryote metabolic profiles that detail community composition across a coastal-offshore biogeochemical gradient obtained using a recently designed autonomous underwater vehicle. The microeukaryote communities were predominantly represented by stramenopiles, dinoflagellates, and ciliates, and were detected throughout the water column, but with the emergence of heterotrophic taxa such as foraminifera, radiolarians, discoba, picozoans, and fungi in deep ocean protein pools (Fig. 7). The metabolic signatures of these surface and deep communities were functionally distinct and indicated viable cells regulating metabolism in response to environmental conditions. We identified several distinctions between transcripts and proteins in taxonomic and functional composition, underscoring the utility of paired metatranscriptomics-metaproteomics for comprehensively reconstructing microbial community dynamics. One of the strongest positive correlations between protein and transcript fractions was a biomarker of nitrogen stress, which was abundant in coastal waters, and supports a tight coupling between nitrogen-related assimilation processes. This multi-omics dataset thus captures shifts in microbial community composition across resource gradients, highlights potential regions of nutrient stress, and offers new insights into the complex relationship between transcripts and proteins in marine microeukaryotes.

**Fig. 7.**
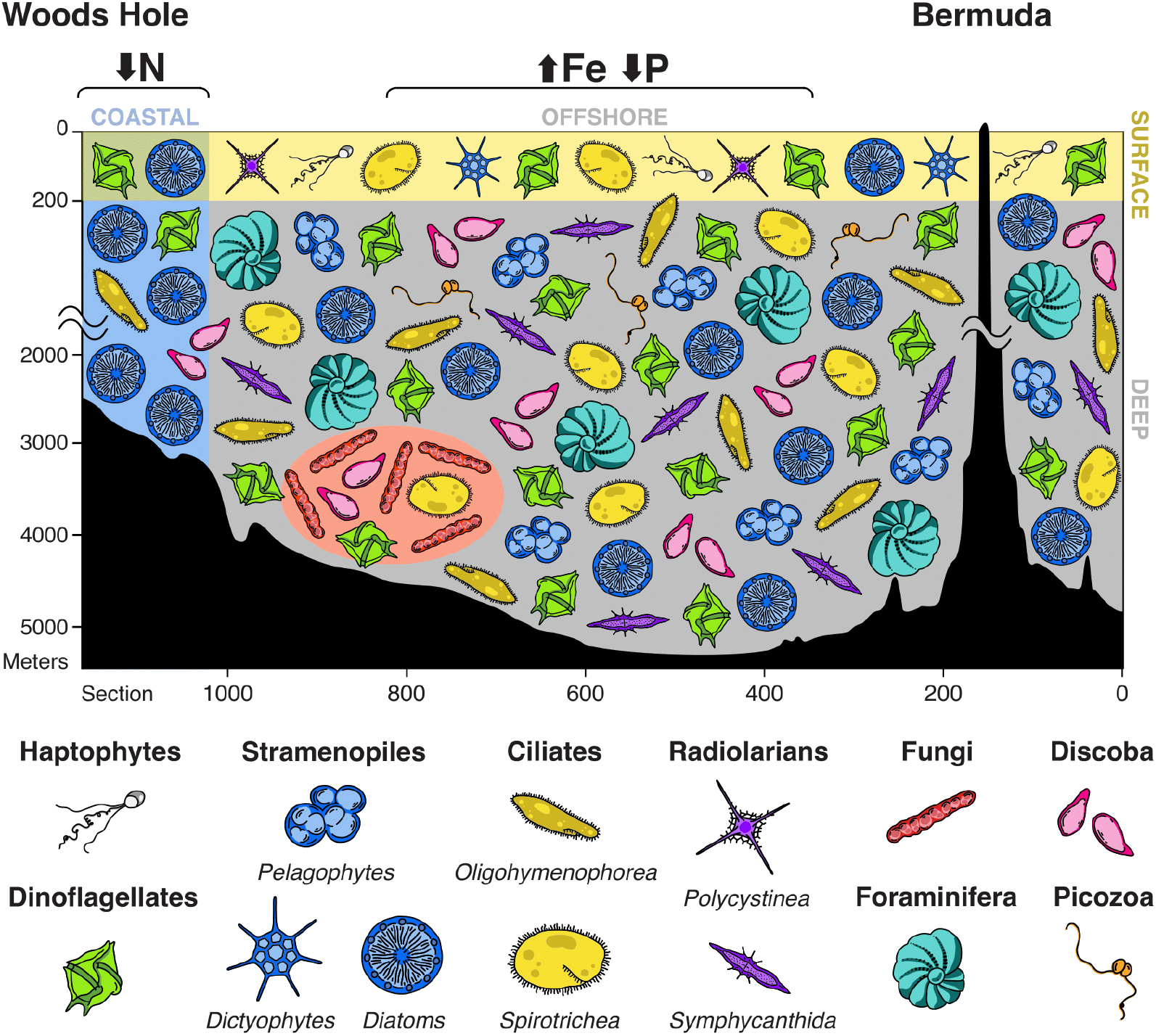
Schematic overview of the western North Atlantic microeukaryote community along the offshore-coastal gradient, integrating protein, transcript and pigment information. Stramenopiles were detected throughout the transect with diatom and dictyophyte proteins in the surface, and diatom and pelagophyte proteins at depth. Diatom pigments were highest in concentration at the coast. Haptophytes proteins were relatively more abundant in surface waters. Dinoflagellate and ciliate proteins and transcripts were detected throughout the water column. Foraminifera, discoba, picozoa and radiolarian proteins were relatively more abundant in the mesopelagic. Radiolarian proteins reflected a vertical community shift with *Polycystinea* in the surface and *Symphycanthida* at depth. Fungi (encapsulated in red circle) appeared in both transcripts and proteins in the bathypelagic. Transcript and proteins indicated nitrate stress in surface coastal waters, with transcriptional signatures of elevated phosphate transport and iron use in offshore waters.

## Methods

### Oceanographic sampling

Oceanographic samples were collected onboard the R/V *Atlantic Explorer* between June 16th - 28th 2019, along a transect beginning at the Bermuda Atlantic Time-series Study (BATS) site and terminating in northeast US continental shelf waters (Woods Hole Oceanographic Institution [WHOI]) (Fig. 1). Biomass was collected using the biogeochemical AUV *Clio* ^27^. *Clio* payloads contained SUPR cartridges housing either 142 mm 0.2 µm polyethersulfone filters (Pall Supor) positioned below 51 µm mesh nylon filters. Alternatively, for pigments, 142 mm combusted GFF filters were used. At Station 1 (BATS), SUPR cartridges with 0.2 µm polyethersulfone filters failed to pump, and biomass was instead collected from *Clio* GFF filters. To complement the vertical collection by *Clio* at this site only, *in situ* battery-operated McLane pumps were used for surface biomass collection (142 mm, 0.2-3 and 3-51 µm). Filters for the biomolecular analysis were sectioned and frozen at −80°C analysis until RNA and protein extractions were performed in the laboratory. GFF filter fractions were additionally preserved for high performance liquid chromatography pigment analyses throughout the surface transect. The R/V *Atlantic Explorer* CTD provided physicochemical contextualization (including temperature, salinity, dissolved oxygen, fluorescence, turbidity used here) and was deployed at all stations except the last one, St. 9, due to time restrictions. Once deployed, *Clio* filtrations happened simultaneously with ship wire operations.

After *Clio* was deployed, trace metal sampling was performed using a trace metal clean rosette equipped with 12 8L X-Niskin bottles on an Amsteel winch line ^87^. Upon retrieval, X-Niskins were brought into a class-100 clean van, and Niskins were pressurized with filtered nitrogen gas. Seawater was filtered through acid-rinsed 142 mm 0.2 µm Supor membranes to separate the particulate and dissolved fractions. Filtered seawater collected into 250 mL HCl-cleaned LDPE bottles were placed in the dark for dissolved metal analyses, and filters for the particulate metal analysis were kept frozen in acid-cleaned cryovials at −80°C until analysis. Subsamples for dissolved nutrients were collected from 0.2 µm-filtered seawater and stored at −20°C. Neither *Clio* nor the trace metal rosette was deployed at St. 4 due to unfavorable weather conditions, although physicochemical data was collected using the ship’s standard CTD.

An Underwater Vision Profiler 5 HD (UVP) ^88^ was deployed with the CTD. The UVP records quantitative, *in situ* images of particles in the water column at a rate of ~15 Hz during CTD descent. Particle images > 0.5 mm Equivalent Spherical Diameter (ESD) are recorded, processed using the Zooprocess software ^89^, identified using the EcoTaxa website ^90^, and validated by a trained taxonomist (full methodology in ^91^). Microeukaryotes were identified to the lowest taxonomic category possible from the images (generally class or order) and abundances were binned by depth, with 40m wide bins from 0-200m, 100m wide from 200-1400m, and 200m wide bins >1400m.

### Metatranscriptomic sequencing & bioinformatic workflow

Forty-four samples across the transect were selected for the metatranscriptomic analysis to achieve coverage along the lateral and vertical ocean gradients. One-sixteenth of 142 mm filters were removed from the −80°C freezer and kept on ice in between handling steps. Total RNA was extracted using the Qiagen RNeasy Mini kit following the quick start protocol with the following modifications. Acid-rinsed 0.1 mm silica beads (Thomas Scientific) were added to each sample along with Qiagen RLT extraction buffer, which was pre-heated to 65°C and contained 2-mercaptoethanol per manufacturer recommendation. Samples were vortexed and physically disrupted in a mini Bead-Beater homogenizer, centrifuged, and filters were removed using forceps that were sterilized with ethanol and RNaseZap (Invitrogen). Lysate was transferred to a Qiashredder column to remove viscosity and improve yields for all but the first set of six randomly chosen samples. Molecular biology grade ethanol (Fisher Scientific) was used to prepare kit buffers for most samples, though the first round of randomly selected six samples were used with reagent grade ethanol and contained impurities. On-column digestion of DNA was performed using DNase I (Qiagen). Elution buffer was pre-heated to 70°C, 30 µL elution volumes were used, and eluate was passed through the column a second time to concentrate RNA.

RNA yields and quality were assessed using a Nanodrop spectrophotometer and Agilent 2100 bioanalyzer. mRNA internal standards were added to each sample for later conversion of sequence read counts to transcript copies ^92^ using poly-adenylated ArrayControl RNA spike #8 (Invitrogen) ^93^ at a standard:sample ng RNA ratio of 0.22%. Since internal standards were not added prior to the extraction, our copies L^−1^ estimates likely overestimate true levels, as nucleic acid loss is expected during the extraction process ^94^. As many samples contained less than 100 ng of total RNA, the SMART-Seq v4 Ultra Low Input RNA Kit (Clontech) for cDNA synthesis was used with 250 pg of RNA input, followed by the NexteraXT DNA library prep kit (Illumina). Libraries were sequenced on the Illumina NovaSeq 6000 platform (2×150 bp PE). Library preparation and sequencing was performed at the Columbia Genome Center.

Sequences were processed using the *euk*rhythmic eukaryotic metatranscriptomic workflow ^95^. This workflow enabled assemblies, read alignments, and open reading frame predictions in an automated, scalable and reproducible Snakemake framework. Briefly, poor quality bases were trimmed using trimmomatic ^96^, and spiked mRNA internal standard reads were removed using bbmap (sourceforge.net/projects/bbmap/). Four different transcriptome assemblers were used on each sample individually: Trinity ^97^, MEGAHIT ^98^, velvet ^99^ and Transabyss ^100^. Assembled contigs were clustered at 100% alignment coverage with cd-hit ^101^ to remove redundant contigs. The assemblies created using the four assemblers were merged by sample, and de-replicated with cd-hit (100% alignment coverage). Next, all 44 samples were merged together using MMseqs2 ^102^ to produce a single dataset-wide final merged assembly. Open reading frames (ORFs) were predicted using TransDecoder (github.com/TransDecoder), retaining nucleotide coding sequences with a minimum length of 300 base pairs. Reads were aligned to both the full-length assembly contigs and the ORF coding nucleotide sequences using Salmon ^103^, which produced pseudocounts and community-wide normalized Transcripts Per Million (TPM) counts, and allowed for a relative comparison across samples. Since the ORFs only include coding regions and sequences longer than 100 AA (300 bp), the mapping percentage is approximately half of that obtained with the full (non-coding) contigs (Supplemental Table 1). In addition to relative analysis, pseudo-read counts were converted to copies L^−1^ following ^92^, with the number of internal standard transcript copies added to samples being approximately 500,000. Taxonomic assessments were performed on ORFs to be consistent with the metaproteomic analysis in which only ORFs (encoded proteins) are considered, as opposed to full-length contigs including non-coding sequences. Functional annotations were performed using eggNOG-mapper ^104^ with default settings.

The ORFs were taxonomically annotated by testing four sequence databases that differed in the reference organisms included: MMETSP + MarRef ^105–107^, EukZoo (github.com/zxl124/EukZoo-database), EukProt ^108^, and PhyloDB (allenlab.ucsd.edu/data). The taxonomic annotations were performed using *EUKulele* with a last common ancestor approach (LCA) ^107^. The LCA conservatively annotated ORFs only when bitscores associated within the top 3% of hits were in taxonomic agreement. If lineage conflicts were present, ORFs were labeled as “NA” (not annotated). In preliminary tests with taxonomic databases, it was determined the MMETSP + MarRef database was missing ecologically relevant protistan and bacterial sequences contained by PhyloDB, EukZoo, and/or EukProt, including *Polycystinea* radiolarians, known inhabitants of Sargasso Sea deep water ^25,26^, and *Spirotrichea* ciliates, which were both relatively abundant in our dataset. We found that prokaryotic ORFs were being erroneously assigned to eukaryotes since they represented the closest available match in the database exceeding quality thresholds (E-value < 10^−5^), as determined using the NCBI non-redundant protein sequence (nr) database which includes a greater number of prokaryotic references. A custom protein database was therefore built using PhyloDB (25,996 taxonomic references, version 1.075), NCBI’s RefSeq (117,030 references, Release 211, ftp.ncbi.nih.gov/refseq/release/complete), two key EukZoo references, and the recently released EukProt database (993 references, version 3) which contains transcriptomes and genomes of diverse eukaryotic lineages ^108^. This combined protein database was not de-replicated, and manual efforts were made to best match coarse classifications belonging to major eukaryotic supergroups at the chosen taxonomic annotation levels analyzed here. Future annotation efforts may benefit from using the entire nr NCBI database which additionally includes incomplete genomes and marine environmental sequences. After top alignment hits within 3% of the highest bit score were identified, taxonomic cutoff levels were called following default *EUKulele* settings using 7 taxonomic levels. At least 15% sequence identity was needed to met at the domain level, 20% at supergroup, 30% at division, 50% at class, 65% at order, 80% at family, and 95% at genus, with 75% consistency among hits needed to make a given taxonomic cutoff ^107^.

### Metaproteomic analysis

One half of the 142 mm filters (0.2-51 μm) were processed for metaproteomics. At St. 1 (BATS), *Clio* failed to recover biomass from the surface layer due to the SUPR cartridges being misaligned, and McLane pumps were used instead with separate 0.2-3 μm (“0.2”) and 3-51 μm (“3”) filter fractions. At this site, additional protein biomass from 20, 175, 250 and 350 m was obtained from *Clio* combusted GFF filters, which were successful in sample collection. Total seawater volumes filtered by *Clio* varied between 30-350 L during this expedition, with generally less volume in surface waters and more at depth due less biomass in deeper layers of the ocean (Supplemental Table 3). The full protein extraction procedure is described in ^37,87^. Briefly, proteins were extracted in an 1% SDS-based detergent in 50 mM HEPES at pH 8.5, reduced with dithiothreitol, alkylated with iodoacetamide, and purified using a polyacrylamide electrophoresis tube gel method. Protein quantification was performed using a BSA assay (BioRad). Trypsin was added to the protein-bead mixture in a 1:20 trypsin:protein ratio. Peptides were purified using C18 tips and diluted to a concentration of 0.1 μg μL^−1^.

Approximately 2-5 µg of purified peptides were injected onto a Dionex UltiMate 3000 RSLCnano LC system with an additional RSLCnano pump, run in online 2D active modulation mode interfaced with a Thermo Fusion mass spectrometer ^109^. The mass spectrometer acquired MS1 scans from 380 to 1,580 m/z at 240K resolution in the Orbitrap. MS2 were collected in data dependent mode in the ion trap with a cycle time of 2 seconds between scans and acquisition of charge states 2 to 10. MS2 scans had 1.6 m/z isolation window, 50 ms maximum injection time and 5 s dynamic exclusion time.

The metatranscriptomic ORFs were used as the protein database, and peptide-spectrum matches were performed using Sequest algorithm within IseNode Proteome Discoverer 2.2.0.388 (Thermo Fisher Scientific) with a parent ion tolerance of 10 ppm and fragment tolerance of 0.6 Da, and 0 max missed cleavage. Identification criteria consisted of a peptide threshold of 98% (false discovery rate [FDR] = 0.1%) and protein threshold of 99% (1 peptide minimum, FDR = 1.5%) in Scaffold 5.1.2 (Proteome Software) resulting in 77,438 proteins and 3,155,061 exclusive spectral counts.

Using the default Scaffold protein inference method (experiment-wide grouping with binary peptide-protein weights), proteins that shared all of their detected tryptic peptides were grouped together and considered the same protein reference (a “protein group”), with the ORF ID randomly chosen among the proteins. Approximately 26% of the ORFs with protein spectral counts were represented by a protein group. Incidents of shared tryptic peptides across distinct taxonomic lineages are generally rare based on investigations in marine prokaryotes ^57^. In cases where a given peptide matched more than 1 protein or protein group, peptides were assigned to the protein with the highest spectral counts. Scaffold’s “exclusive spectral counts” were used for the proteomic analysis to avoid double counting of spectral counts, i.e. assigning counts to more than one protein/protein group ^37,110^. Protein spectral count concentrations per volume seawater were calculated using equation (3), adapted from ^110^:

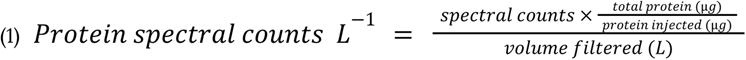

### Macronutrients & pigments

Dissolved nitrate+nitrite, nitrite and silicate were measured on an Alpkem Rapid Flow Analyzer, and ammonium and phosphate were measured on a Technicon AutoAnalyzer II at Oregon State University. For most stations, nutrients were collected and measured from trace metal rosette niskin bottles. However, St. 8 nutrients showed oceanographically inconsistent profiles, potentially due to samples thawing in transit. Nutrients collected from *Clio* at St. 8 were used instead, with general agreement between trace metal rosette and *Clio*-collected nutrients at other stations and depths.

Pigments were collected onto 142 mm GFF filters by *Clio* at the same depths that proteins were collected on Supor filters. GFF 25 mm punchouts were used for the pigment analysis, and the estimated volume filtered through 25 mm was used to quantify pigments per volume. Concentrations are reported as ng/kg assuming the standard density of seawater (1025 kg/m^3^). At stations 1 and 3, GFF fractions for pigments were not collected by *Clio*. Instead, pigments were analyzed from the CTD rosette at these stations whereby samples (4 L) were filtered directly onto 25 mm GFF glass fiber filters and immediately frozen in liquid nitrogen. Samples were analyzed on an Agilent 1100 series high performance liquid chromatography (HPLC) using the method of ^111^ and instrument calibration was performed with certified pigment standards from the Danish Hydraulic Institute.

### Dissolved trace metals

Filtered seawater was acidified to pH 1.8 using hydrochloric acid (Optima grade, Fisher Chemical), and samples intended for dissolved trace metal analyses were stored for 6 months at room temperature in the dark before metal quantification. Seawater preconcentration was performed using the seaFAST automated preconcentration system ^112^ followed by quantification via inductively coupled plasma mass spectrometry. Reagents consisted of a 4M ammonium acetate pH 6.0 buffer prepared using high purity ammonium hydroxide and acetic acid (Optima grade, Fisher Chemical), a 1% nitric acid rinse solution (Optima grade, Fisher Chemical), 10% nitric acid elution acid, and a second internal standard 10% nitric elution acid solution containing 10 ppb indium (^115^In) (SPEX CertiPrep). All solutions were prepared with 18.2 Ω Milli-Q purified water (Millipore). Polypropylene conical tubes used with the autosampler were HCl acid-soaked and pH 2-rinsed prior to use. Process blanks consisted of Milli-Q HCl-acidified to pH 2 (Optima grade, Fisher Chemical), and were run alongside samples. A stable isotope cocktail, which consisted of ^57^Fe, ^61^Ni, ^65^Cu, ^67^Zn and ^111^Cd, was spiked (50 µL) into each 15 mL sample to account for recovery and matrix effects. Stable isotope spike standards were prepared by dissolving metal oxides into 50% nitric acid and diluted using volumetric flasks (Optima grade, Fisher Chemical). Spike concentrations were chosen based on the amount needed to generate an approximate 1:1 ratio of spiked:stable isotope in the 15 mL sample ^113,114^.

Following offline seaFAST preconcentration to 500 μL, the samples were analyzed using an iCAP Q inductively coupled plasma-mass spectrometer (ICP-MS) (Thermo Scientific). An external standard curve was used with multi-element and indium (In) standards (SPEX CertiPrep), diluted to range from 1-10 ppb in 5% nitric acid. Dissolved metal concentrations were determined using the following isotope dilution equation (1):

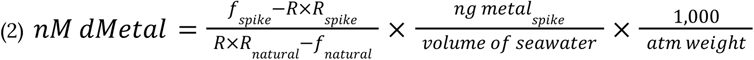

where, using Fe for example, *R* = ratio of ^56^Fe/^57^Fe in sample, *R_spike_* = ^57^Fe_cps_ / (^56^Fe_cps_ + ^57^Fe_cps_) in the spike, *R_natural_* = ^57^Fe_cps_ / (^56^Fe_cps_ + ^57^Fe_cps_) in 10 ppb standard, *atm weight* = average atomic weight between the two isotopes, *f_spike_* = ^56^Fe_cps_ / (^56^Fe_cps_ + ^57^Fe_cps_) in spike, *f_natural_* = ^56^Fe_cps_ / (^56^Fe_cps_ + ^57^Fe_cps_) in 10 ppb standard, and *ng spike* = the metal concentration in the spike, determined empirically using ^57^Fe_cps_ of spike, and manually tuned to the value that allowed for the 10 ppb + spike standard to equal 5 ng. As there is no second stable isotope for manganese (Mn), concentrations were determined using the ^57^Fe isotope spike.

Nitric acid (5%) injection blanks were subtracted from sample metal cps values except for Zn, in which injection blanks were generally higher than process blanks. Accuracy was determined using the 2009 GEOTRACES surface coastal (GSC) seawater intercalibration standard (n=5): dFe = 1.65 ± 0.19 nM [GSC = 1.56 ± 0.12 nM], dZn = 1.65 ± 0.17 nM [GSC = 1.45 ± 0.10 nM], dCu = 1.38 ± 0.16 nM [GSC = 1.12 ± 0.15 nM], dCd = 0.37 ± 0.03 nM [GSC = 0.37 ± 0.02 nM], dNi = 4.28 ± 0.17 nM [GSC = 4.5 ± 0.21 nM], and dMn = 2.42 ± 0.29 nM [GSC = 2.23 ± 0.08 nM].

The limit of detection was determined by calculating 3x the standard deviation of process blanks dataset-wide: dFe = 0.23 nM (n=14), dZn = 0.35 nM (n=16), dCu = 0.03 nM (n=18), dCd = 0.0103 nM (n=16), dNi = 0.07 nM (n=18), dMn = 0.03 nM (n=18). Blanks that were overtly contaminated with Fe (4 of 18), Zn (2 of 18) or Cd (2 of 18) were not included in the LOD estimation. In the case of Zn, high deep water (>1,000 m) concentrations altered the spiked:stable isotope ratio, and accurate concentrations were not able to be obtained. Measured concentrations below the LOD are reported as “below detection” in trace metal profiles, and “0” in section plots and correlation matrices. Station 5 surface concentrations have been published as part of dissolved organic matter photochemistry and carbon monoxide cycling experiments in the Sargasso Sea (Zafiriou et al. submitted), and the full dissolved metal dataset has been uploaded to BCO-DMO under project 765945.

### Statistical approaches, normalizations & visualization

Normalizations were performed at the community-wide level for ordinations, network analyses, and the partial least squares (PLS) regression analyses. For these community-wide analyses, transcripts were normalized following the transcripts per million approach, which enables relative comparison of gene expression across samples by accounting for differences in library sizes and the length of ORFs. Proteomic exclusive spectral counts were normalized using the normalized spectral abundance factor (NSAF) calculation, which follows the TPM approach.

Non-metric multidimensional scaling (NMDS) ordinations were performed using community-wide normalized transcripts and protein counts associated with ORFs having both a taxonomic and functional annotation (eukaryotic ORFs assigned by EUKulele and KEGG Orthology ID assigned by eggNOG-mapper [^115^]). Counts were summed to the supergroup level and log_2_ normalized, and Bray-Curtis used to calculate distance using the *metaMDS* function in *vegan*.

Undirected networks were built to separately identify co-occurring eukaryotic transcripts or proteins. To reduce the size of the networks, only ORFs with a taxonomic and functional annotation were used, and counts were summed to the supergroup and KEGG annotation level (e.g., Chlorophyta_ko:K00053). Annotated ORFs with community-wide TPM >10 or NSAF counts >0.01 were retained for the analysis, and counts were log_2_ transformed. A correlation matrix was calculated using Spearman rank correlation distances in *igraph* ^116^. Correlations >|0.65| were considered. The graph was converted to a minimum spanning tree (MST) to identify the minimal number of edges needed for all nodes to be connected, thus decreasing the size of the network. Small, unconnected subnetworks were removed. Clusters of correlated genes (modules) were identified using fast greedy modularity optimization (“cluster fast greedy”) and the networks were visualized with the Kamada-Kawai algorithm. Modules were investigated through taxonomic and functional composition. KEGG IDs were grouped into BRITE categories for a broader functional classification, minus the “Enzyme” category as this grouping was considered too broad to be biologically meaningful. Environmental context for the modules was determined by calculating eigengene values, or the first principle component of gene/protein expression representing each module using *tmod*, and calculating Spearman rank correlations with environmental data. These correlations were only calculated for stations/depths where corresponding trace metal data was collected.

Partial least squares (PLS) regression was performed to correlate protein abundance and gene expression. Only eukaryotic ORFs with a taxonomic and functional annotation were used, and counts were summed across eukaryotic taxa to the functional level. Annotated ORFs with community-wide TPM >5 or NSAF counts >0.01 were retained for the analysis. KEGG Orthology carbon and nutrient transporters largely compiled by ^117^ were identified in the dataset, excluding the following proteins: an archaeal phosphate transport substrate-binding protein (pstS; K02040) and bacterial iron complex outer membrane receptor protein (TC.FEV.OM; K02014). ORF sequences annotated as these proteins showed high percent identity to eukaryotes in our early comprehensive taxonomic database (PhyloDB + RefSeq), but upon further examination, were closer matches to prokaryotes contained within the NCBI non-redundant protein database, which includes other relevant prokaryotic references as well as marine metagenome-assembled genomes. The PLS regression was performed using *mixOmics* with default scaling and using two components, and was evaluated using leave one out cross validation ^118^.

## Data Availability

AE1913 CTD data and processed trace metal, and macronutrient concentrations are publicly available on BCO-DMO (https://www.bco-dmo.org/project/765945). Raw sequences have been deposited to the NCBI SRA database under BioProject ID PRJNA903389 (https://www.ncbi.nlm.nih.gov/bioproject/PRJNA903389). Metatranscriptomic assembly, annotations and count data are available through Zenodo (https://zenodo.org/record/8287779). Metaproteomic mass spectrometry data is available on ProteomeXchange through PRIDE accession PXD045395. Analysis notebooks are available on GitHub (https://github.com/cnatalie/BATS).

## Acknowledgements

We are grateful to the captain, crew, and science party of the AE1913 BATS cruise onboard the *R/V Atlantic Explorer*. The Clio deployments and recoveries were made possible by the expertise of the BIOS team. We thank members of the Seth John Lab (USC), especially Rachel Kelly and Xiaopeng Bian, for guidance and useful discussions regarding the isotope dilution calculations. We thank Maggi Mars Brisbin for valuable proteomic comments and discussion. The NIH/NCI Cancer Center Support Grant P30CA013696 supported Columbia’s Genomics and High Throughput Screening Shared Resource where metatranscriptomes were sequenced. Bioinformatic processing and computational analyses were performed on Woods Hole Oceanographic Institution’s high performance computing cluster, Poseidon. NRC was supported by Simons Foundation #544236 and the University of Georgia Skidaway Institute of Oceanography, and by the National Science Foundation (NSF) during the time of writing (OPP-2240780 and OPP-2149071). AIK was supported by the DOE Computational Science Graduate Fellowship (DE-SC0020347). JAB was supported by NSF OCE-1333212, OCE-1658067, OCE-1924508 and NOAA #NA16SEC4810009. MAS was supported by Simons Foundation #1038971 and NSF OCE-1658030.

## Supplemental Tables

**Supplemental Table 1**. Total dissolved metal concentrations (nM), macronutrients (μM), and particulate metals (pM) for each station and depth.

**Supplemental Table 2**. Metatranscriptomic assembly statistics, including read mapping percentages and number of annotated open reading frames.

**Supplemental Table 3**. Total protein extracted from filters (μg), amount injected onto the mass spectrometer (μg), and volume filtered (L) by *Clio*.

**Supplemental Table 4**. Taxonomic groups and KEGG functional processes associated with each module of the protein network.

**Supplemental Table 5**. Taxonomic groups and KEGG functional processes associated with each module of the transcript network.

**Supplemental Table 6.** Summary of *Clio* dives during AE1913.

**Supplemental Table 7**. TPM-normalized transcript counts for each eukaryotic open reading frame (ORF). Only ORFs with single KO annotations are included.

**Supplemental Table 8**. NSAF-normalized protein spectral counts for each eukaryotic open reading frame (ORF). Only ORFs with single KO annotations are included.

## Supplemental Figures

**Supplemental Fig. 1.**
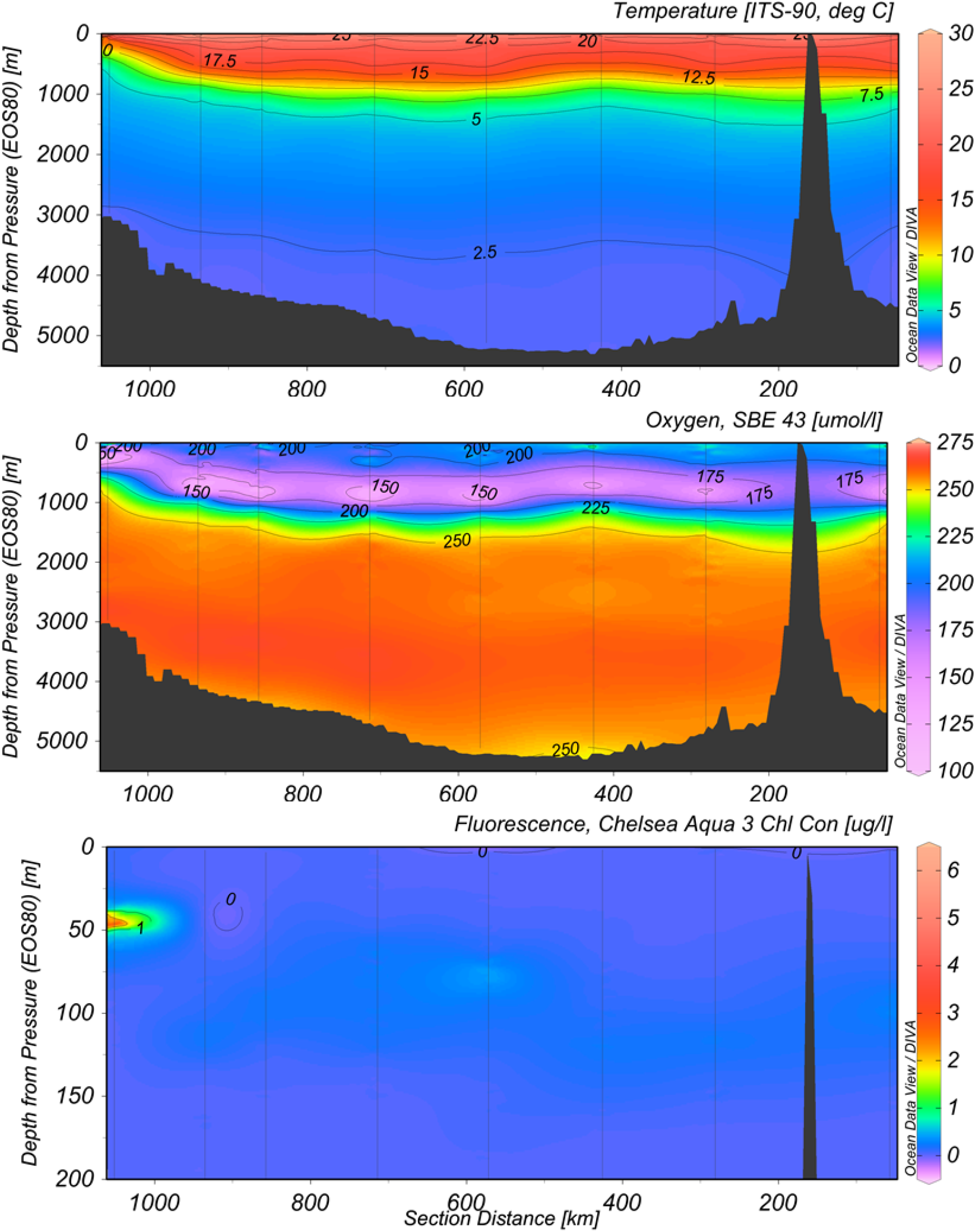
Continuous beam temperature, dissolved oxygen, and fluorescence measurements from the full depth water column, obtained with the R/V *Atlantic Explorer* CTD rosette.

**Supplemental Fig. 2.**
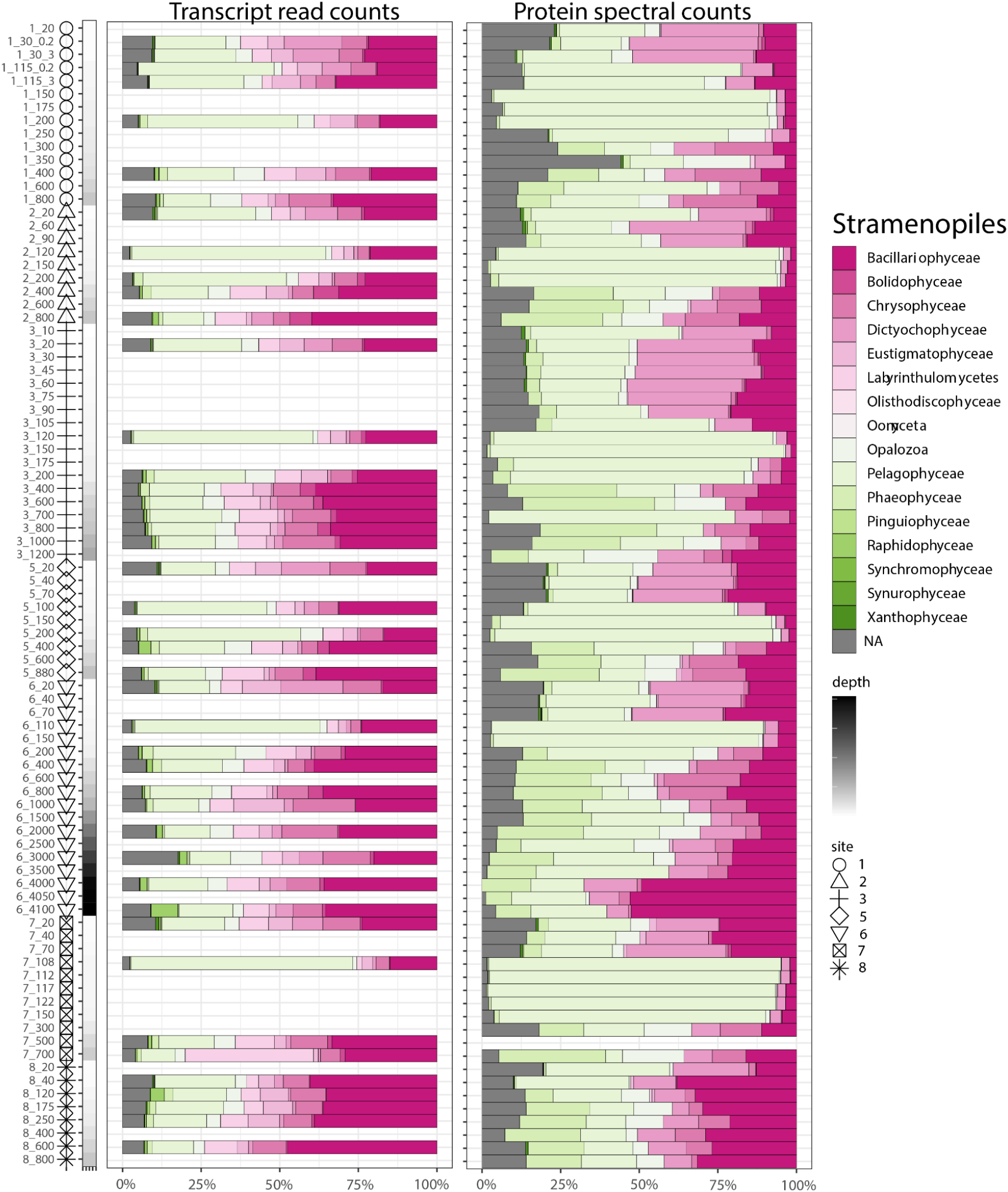
Relative abundance of <u>stramenopile</u> transcript reads recruited to ORF predicted proteins (left), and exclusive proteomic spectral counts assigned to ORFs (right). Each row represents a station and depth (“7_700” = St. 7, 700 m). NA represent haptophytes with lineage conflicted taxonomic assignments unable to be classified at the level shown.

**Supplemental Fig. 3.**
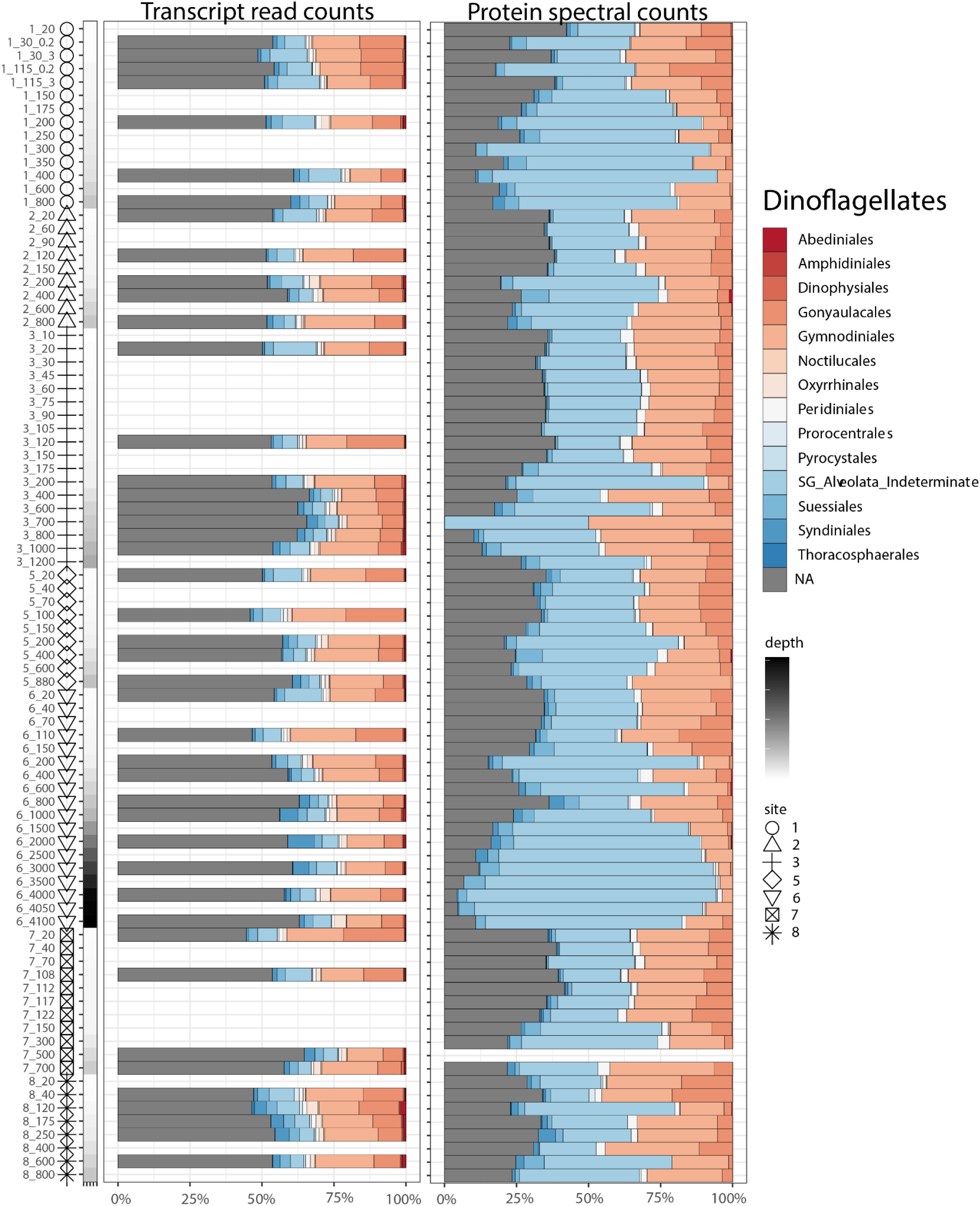
Relative abundance of <u>dinoflagellate</u> transcript reads recruited to ORF predicted proteins (left), and exclusive proteomic spectral counts assigned to ORFs (right). Each row represents a station and depth (“7_700” = St. 7, 700 m). NA represent haptophytes with lineage conflicted taxonomic assignments unable to be classified at the level shown.

**Supplemental Fig. 4.**
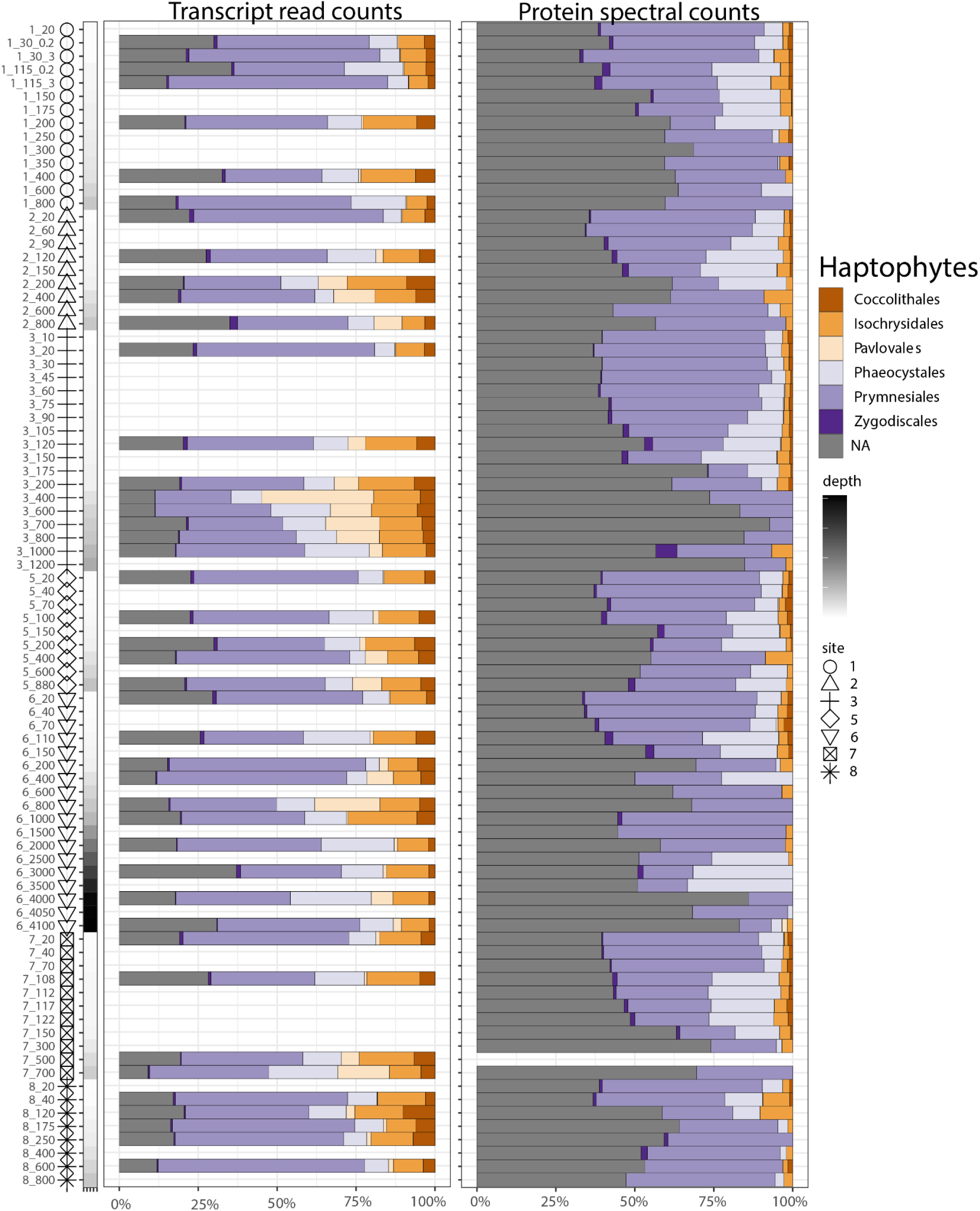
Relative abundance of <u>haptophyte</u> transcript reads recruited to ORF predicted proteins (left), and exclusive proteomic spectral counts assigned to ORFs (right). Each row represents a station and depth (“7_700” = St. 7, 700 m). NA represent haptophytes with lineage conflicted taxonomic assignments unable to be classified at the level shown.

**Supplemental Fig. 5.**
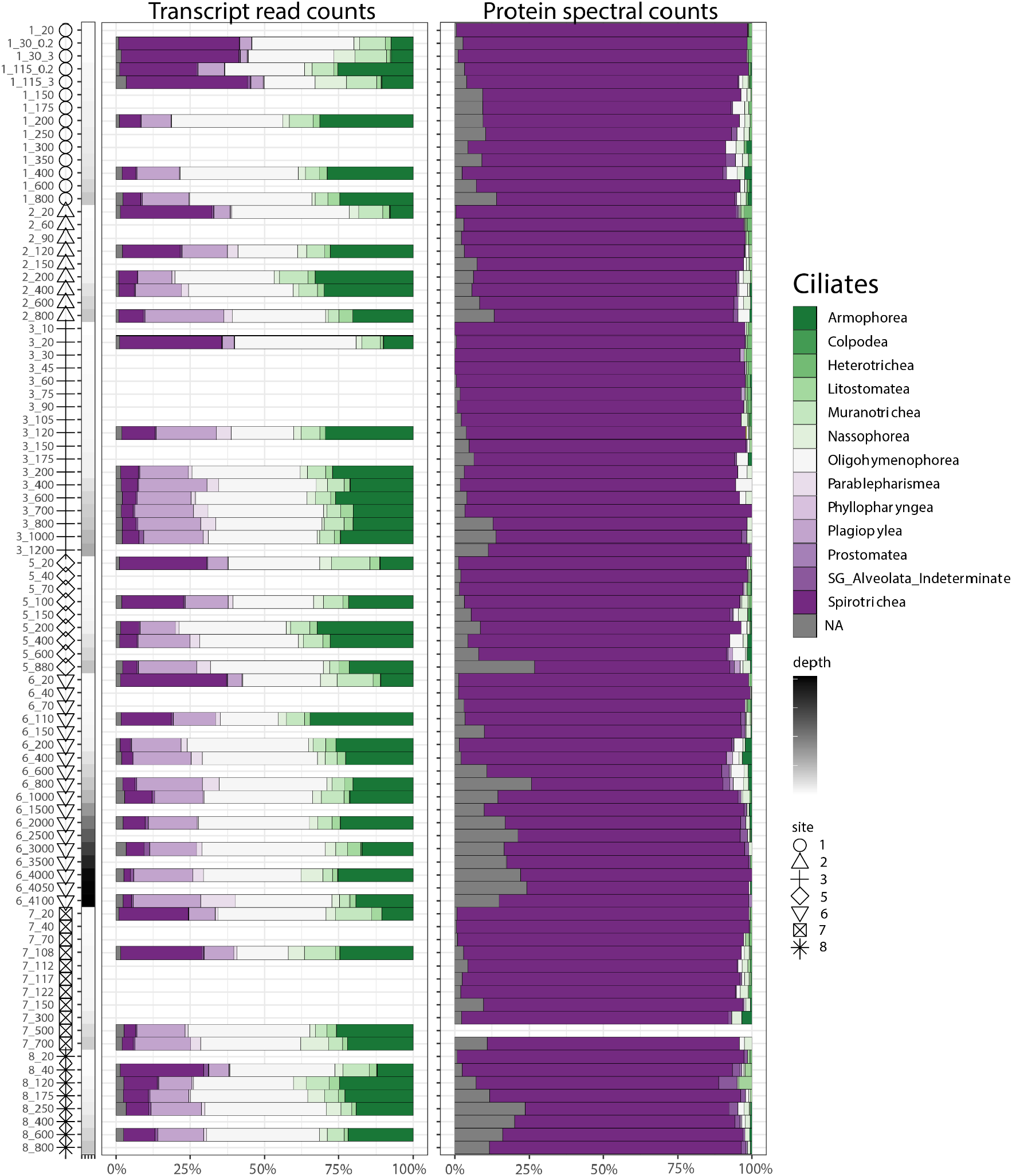
Relative abundance of <u>ciliate</u> transcript reads recruited to ORF predicted proteins (left), and exclusive proteomic spectral counts assigned to ORFs (right). Each row represents a station and depth (“7_700” = St. 7, 700 m). NA represent haptophytes with lineage conflicted taxonomic assignments unable to be classified at the level shown.

**Supplemental Fig. 6.**
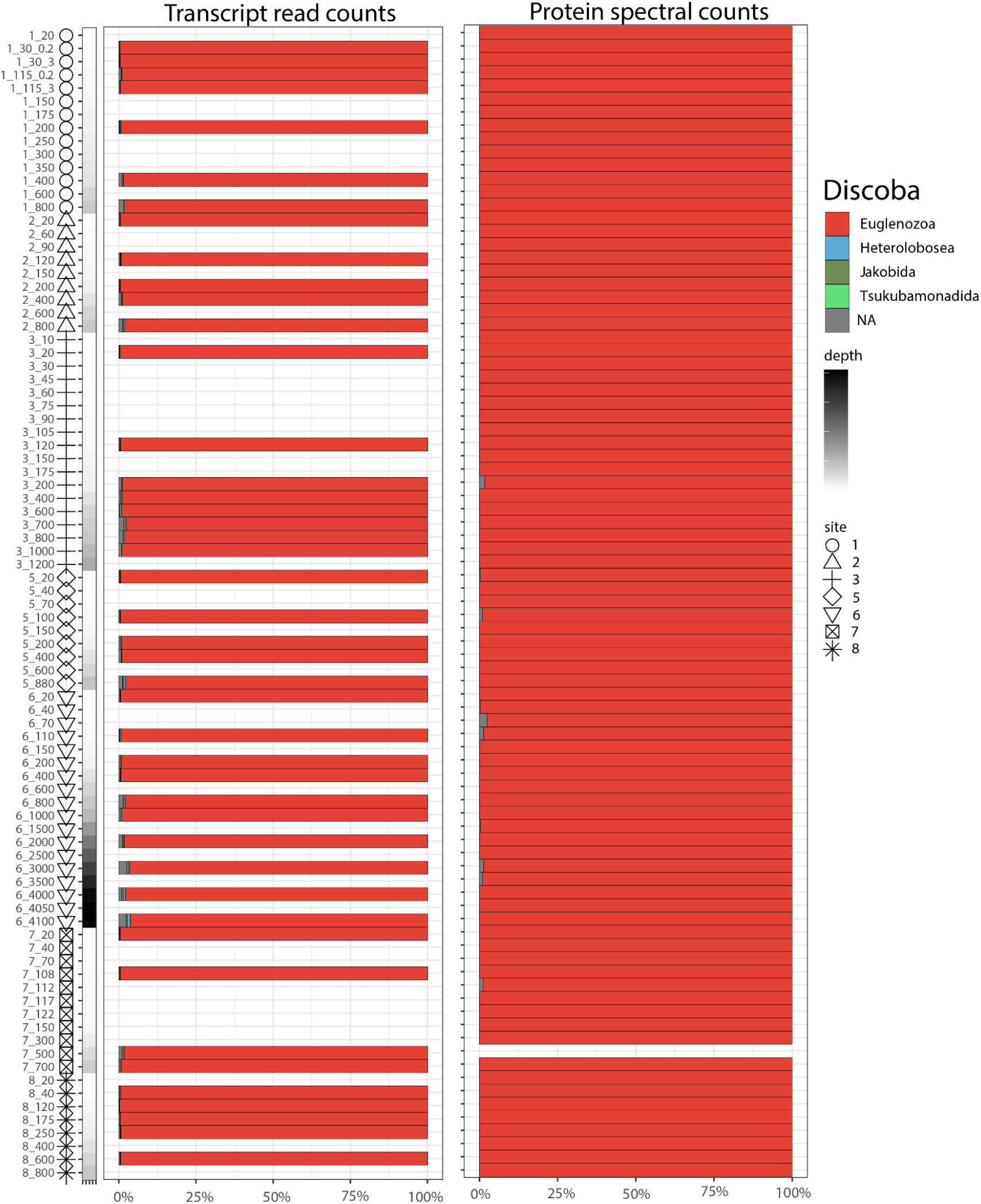
Relative abundance of <u>discoba</u> transcript reads recruited to ORF predicted proteins (left), and exclusive proteomic spectral counts assigned to ORFs (right). Each row represents a station and depth (“7_700” = St. 7, 700 m). NA represent haptophytes with lineage conflicted taxonomic assignments unable to be classified at the level shown.

**Supplemental Fig. 7.**
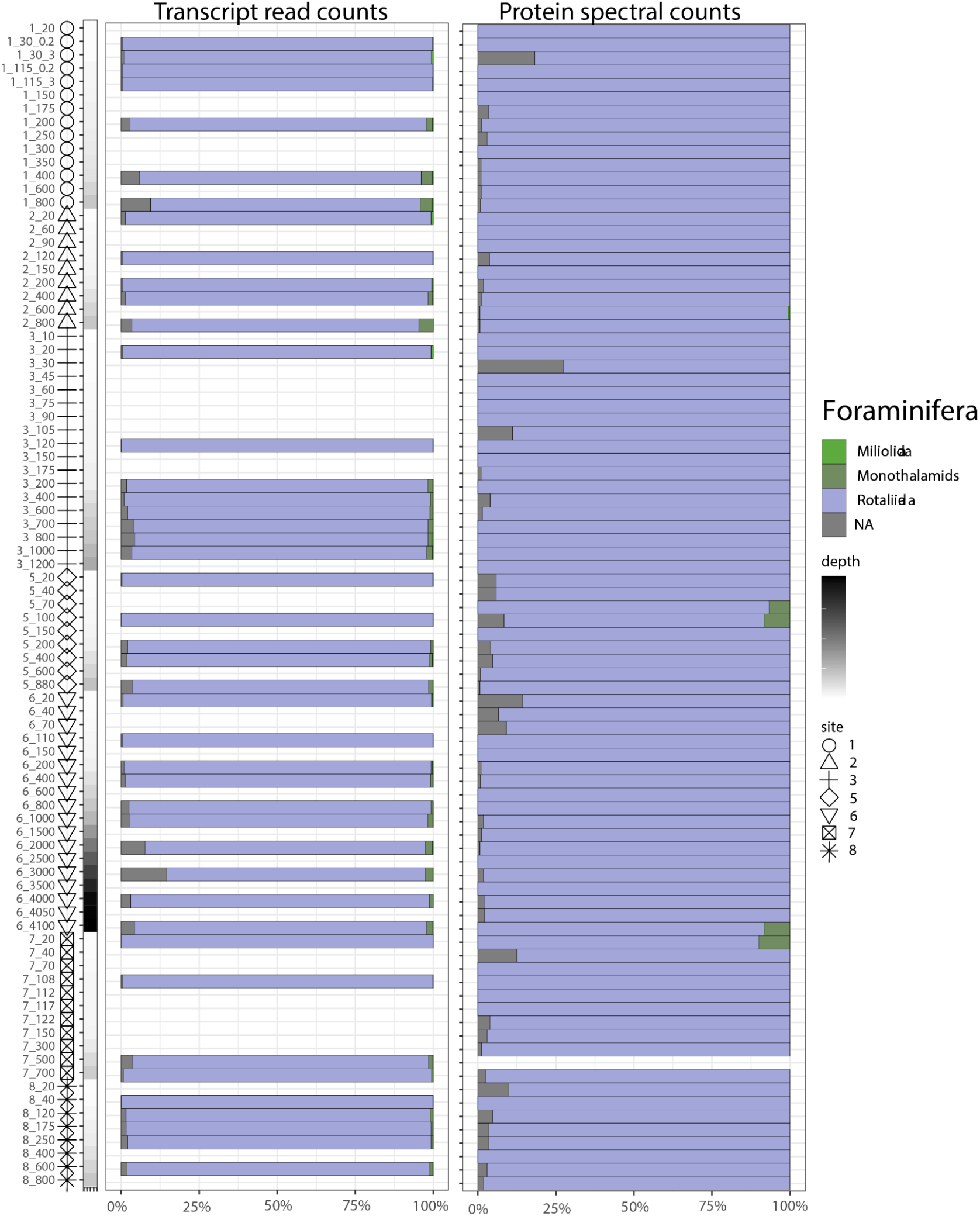
Relative abundance of <u>formaninifera</u> transcript reads recruited to ORF predicted proteins (left), and exclusive proteomic spectral counts assigned to ORFs (right). Each row represents a station and depth (“7_700” = St. 7, 700 m). NA represent haptophytes with lineage conflicted taxonomic assignments unable to be classified at the level shown.

**Supplemental Fig. 8.**
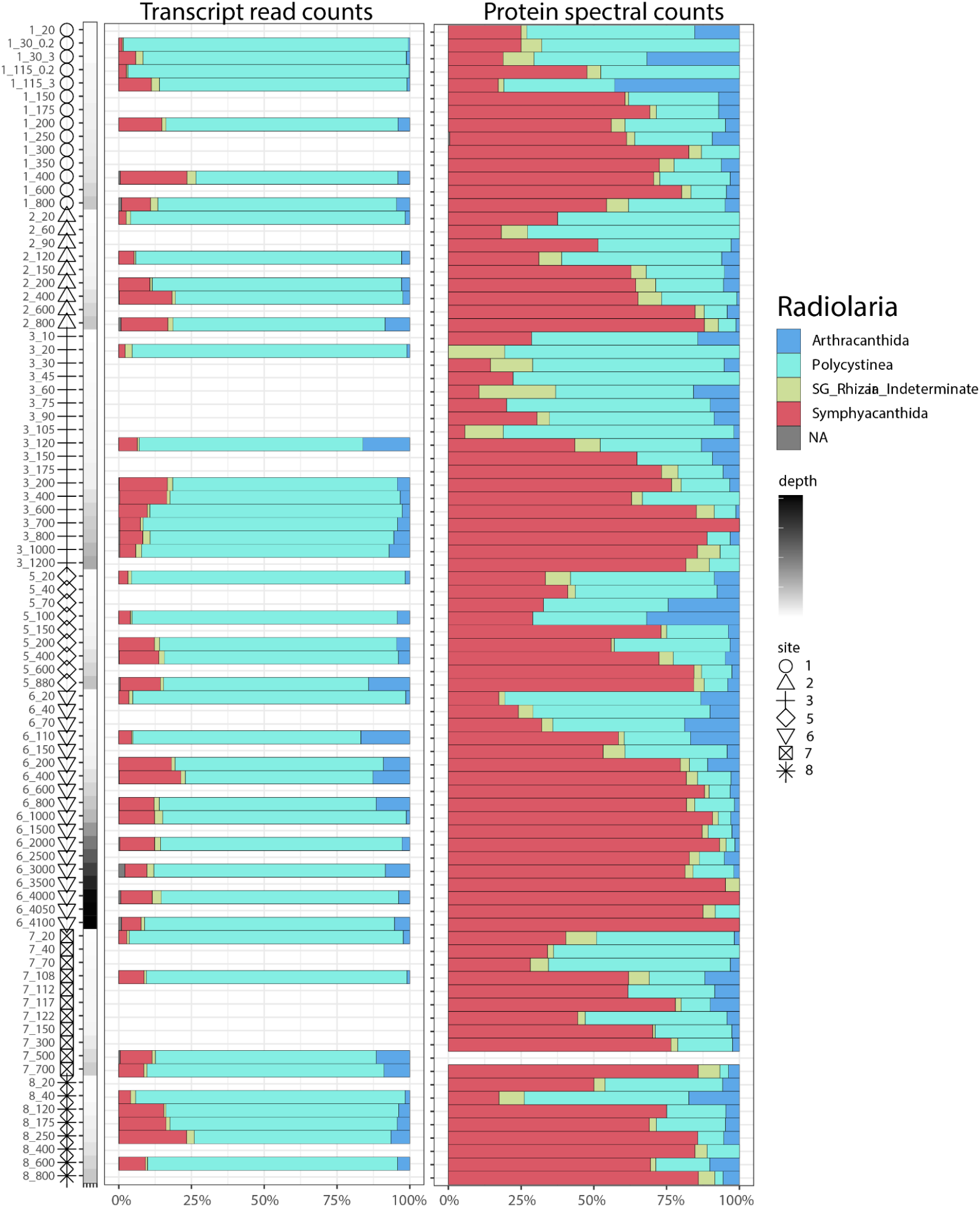
Relative abundance of <u>radiolaria</u> transcript reads recruited to ORF predicted proteins (left), and exclusive proteomic spectral counts assigned to ORFs (right). Each row represents a station and depth (“7_700” = St. 7, 700 m). NA represent haptophytes with lineage conflicted taxonomic assignments unable to be classified at the level shown.

**Supplemental Fig. 9.**
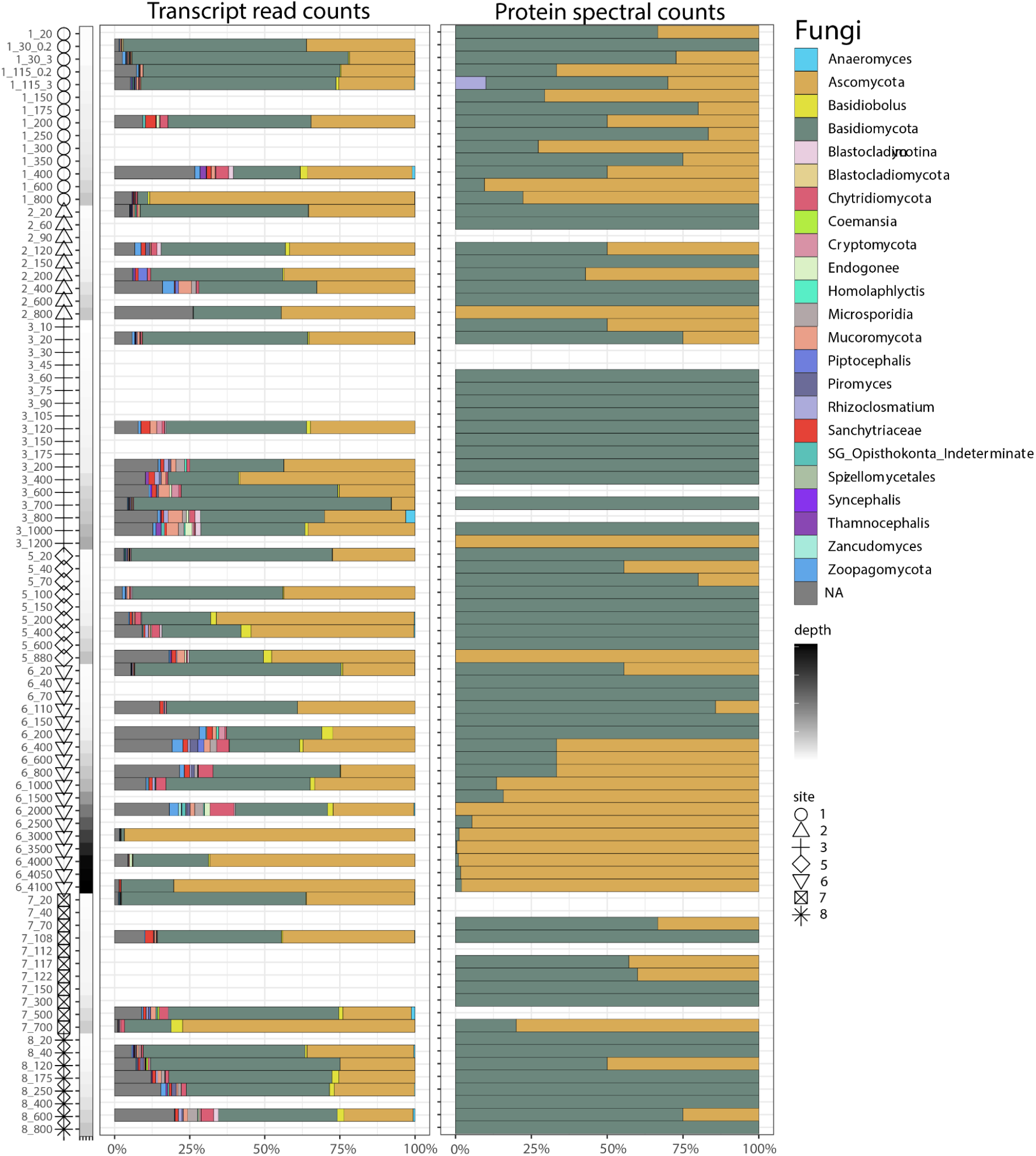
Relative abundance of <u>fungi</u> transcript reads recruited to ORF predicted proteins (left), and exclusive proteomic spectral counts assigned to ORFs (right). Each row represents a station and depth (“7_700” = St. 7, 700 m). NA represent haptophytes with lineage conflicted taxonomic assignments unable to be classified at the level shown.

**Supplemental Fig. 10.**
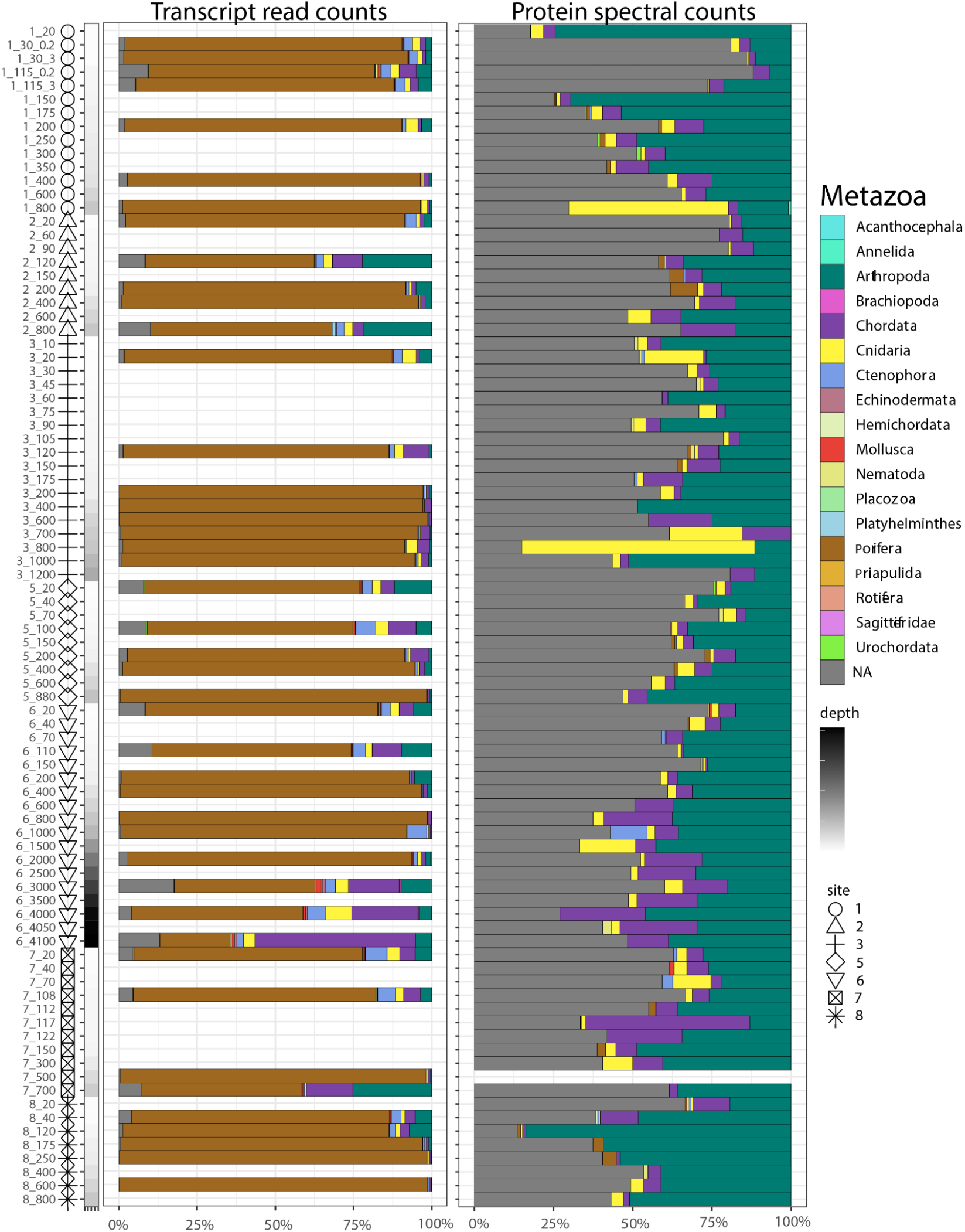
Relative abundance of <u>metazoa</u> transcript reads recruited to ORF predicted proteins (left), and exclusive proteomic spectral counts assigned to ORFs (right). Each row represents a station and depth (“7_700” = St. 7, 700 m). NA represent haptophytes with lineage conflicted taxonomic assignments unable to be classified at the level shown.

**Supplemental Fig. 11.**
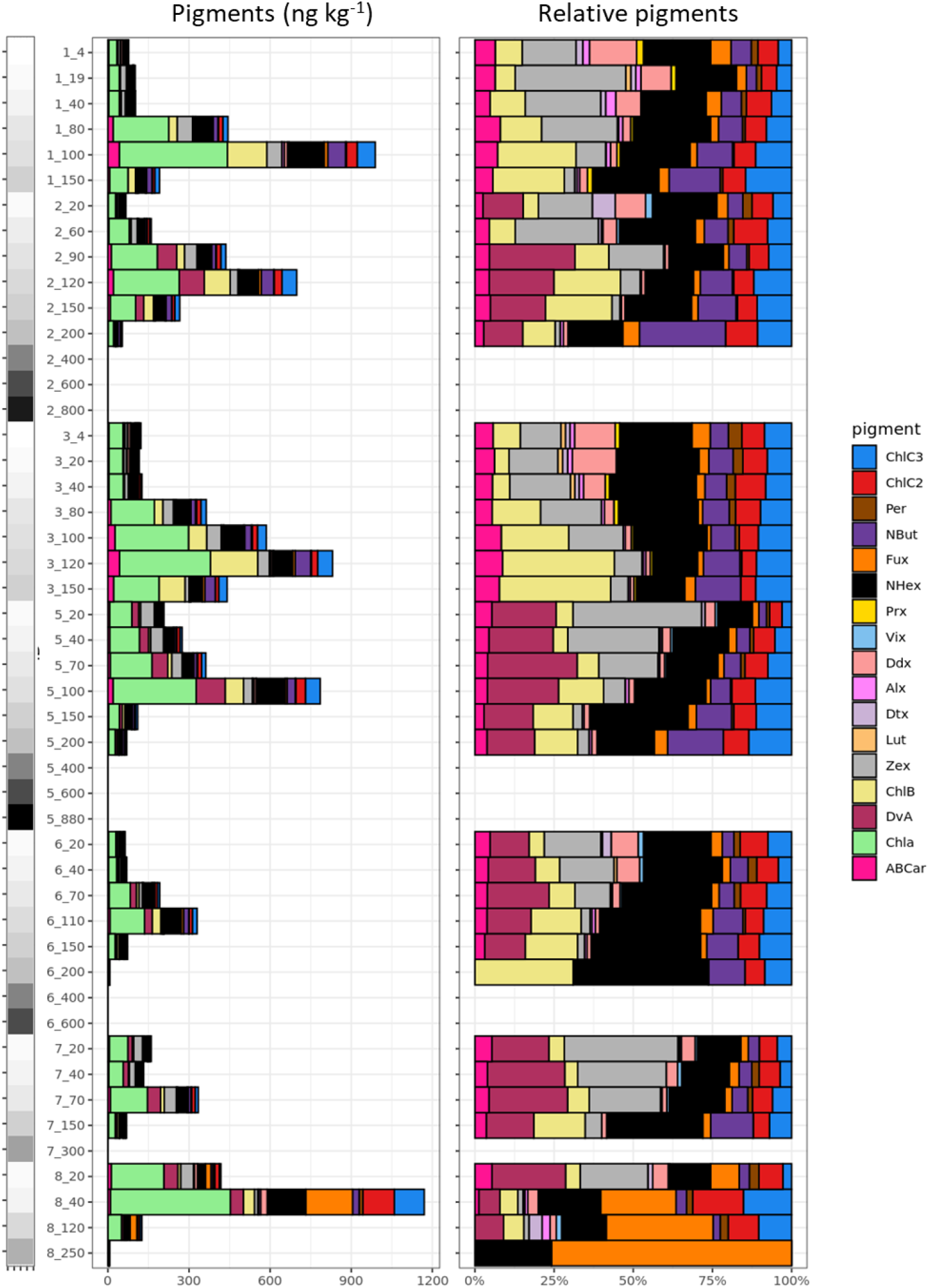
Pigment concentrations (left) and relative pigment composition (right; excluding chlorophyll *a*) across stations and depths. Maximum depths measured for pigments ranged between 150-880 m across the transect. Pigments were not measured from *Clio* dives at Stations 1 or 3, and instead pigments were collected from the CTD casts, which generally agreed with *Clio* measurements at other stations. Note that DvA and Vix were not measured at St. 1 and 3 from the CTD casts. Pigment concentrations below 400 m were undetected. The common pigment chlorophyll *a* is not depicted in the relative abundance plot to highlight other pigment composition. [Pigment abbreviations: ChlC3 = Chlorophyll c3, ChlC2 = Chlorophyll c2, Per = Peridinin, NBut = 19’But-Fucoxanthin, Fux = Fucoxanthin, NHex = 19’Hex-Fucoxanthin, Prx = Prasinoxanthin, Vix = Violoxanthin, Ddx = Diadinoxanthin, Alx = Alloxanthin, Dt = Diatoxanthin, Lut = Lutein, Zex = Zeaxanthin, ChlB = chlorophyll *b*, DvA = Divinyl Chlorophyll a, Chla = chlorophyll *a*, ABCar = Alpha & Beta Carotene.]

**Supplemental Fig. 12.**
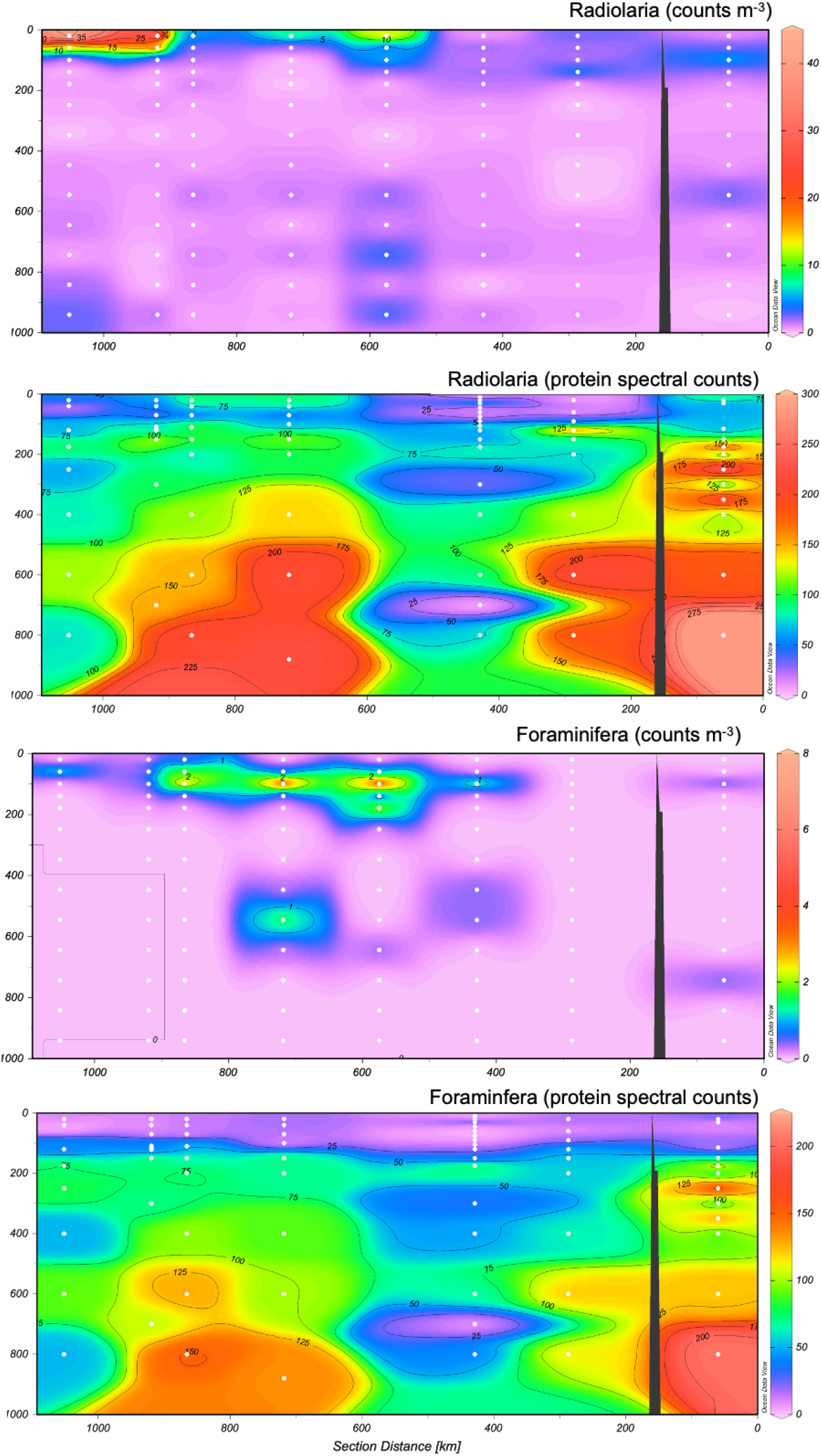
Absolute counts per m^3^ of radiolarians (combined *Collodaria, Phaeodaria, and Acantharea*) and foraminifera >500 μm estimated from Underwater Vision Profiler (UVP) images and total protein spectral counts. UVP counts were binned by depth, with 40m wide bins from 0-200m, 100m wide 200-1400, and 200m wide bins >1400m. Concentrations are visualized for the median depth of each bin. Protein spectral counts are not normalized.

**Supplemental Fig. 13.**
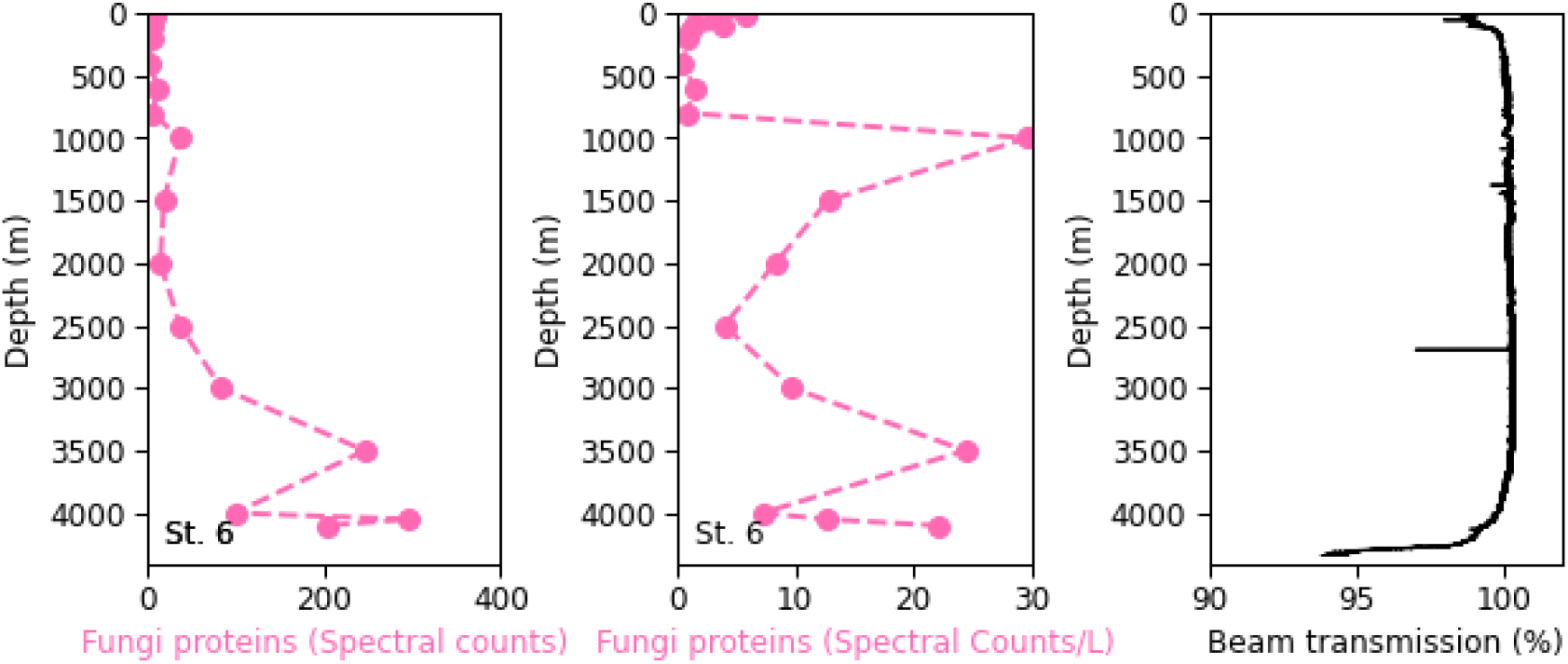
Fungi protein spectral counts (left), fungi proteins normalized for total protein extracted, injected, and seawater filtered (middle), and beam transmission (turbidity; right) at St. 6, where full-depth CTD and protein samples were collected.

**Supplemental Fig. 14.**
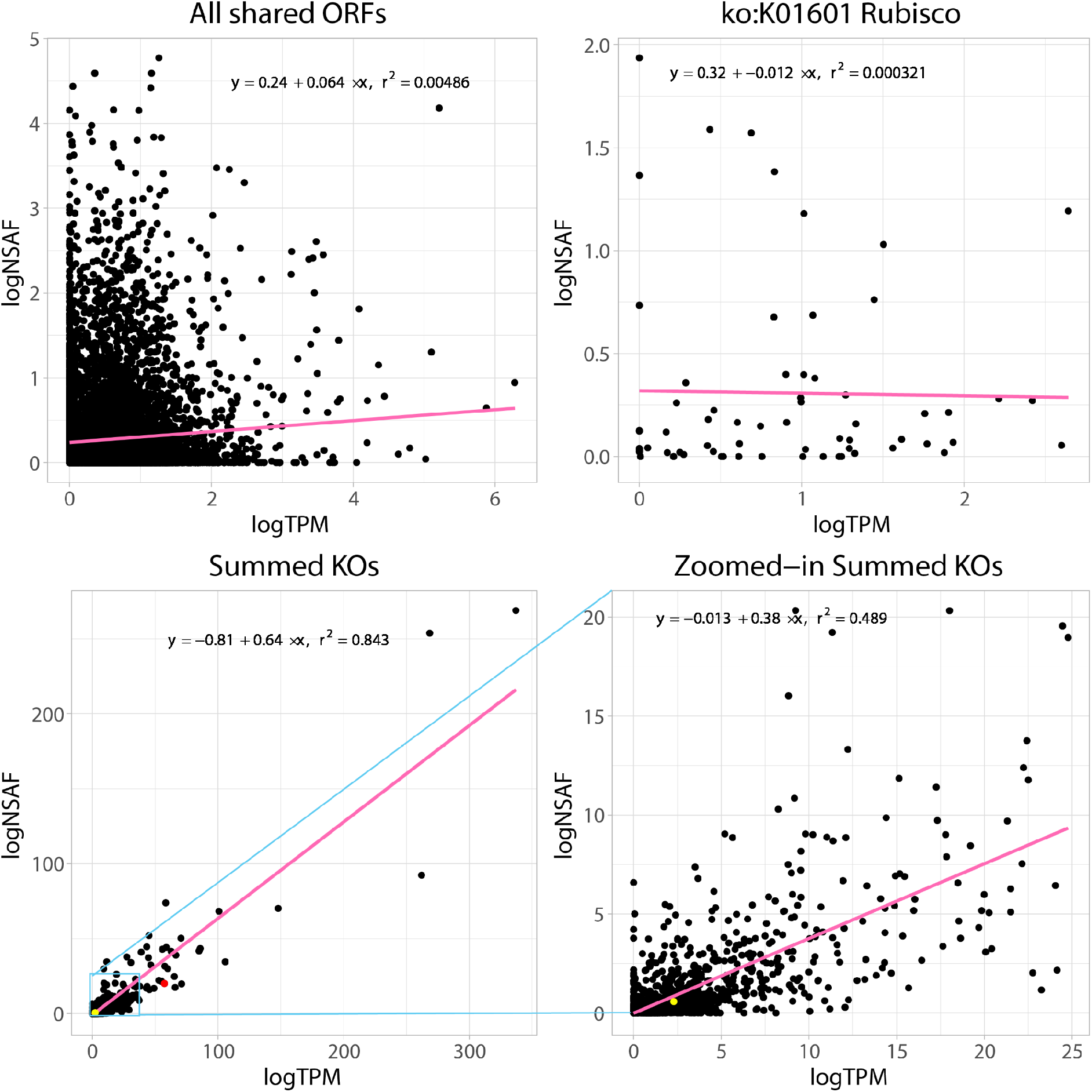
Dataset-wide correlations between eukaryotic transcripts and proteins. Counts were summed across all sites/depths and log-transformed. (Only functionally annotated ORFs with single KO annotations are included. (**A**) Correlations on the open reading frame (ORF) level show weak agreement. (**B**) Subset ORFs including only Rubisco (KEGG Orthology K01601) likewise show a weak correlation between transcripts and proteins. (**C**) ORFs summed to functional processes show stronger agreement between pools, including those in low overall abundance (**D**). Yellow = Rubisco (K01601). Red = Nitrate transporter NRT (K02575).

**Supplemental Fig. 15.**
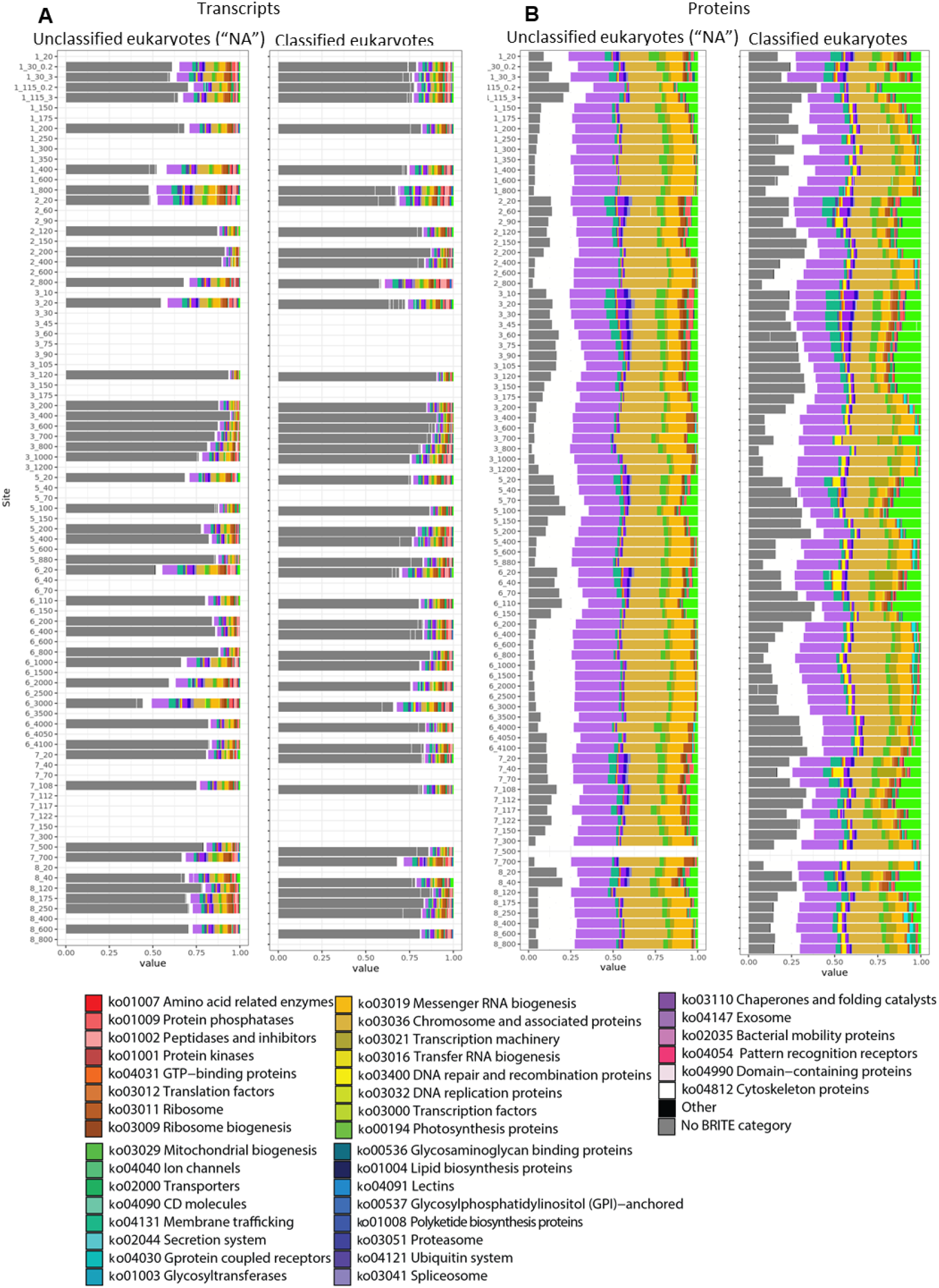
Functional composition of eukaryotic ORFs with a taxonomic assignment (classified eukaryotes) compared to ORFs with conflicts across eukaryotic supergroups (unclassified eukaryotes [“NA”]), in the transcript (**A**) and protein fractions (**B**). Only ORFs with an *eggnog-mapper* annotation are shown, with Kegg BRITE categories listed by color. Gray represents *eggnog-mapper* annotations without a Kegg BRITE classification.

**Supplemental Fig. 16.**
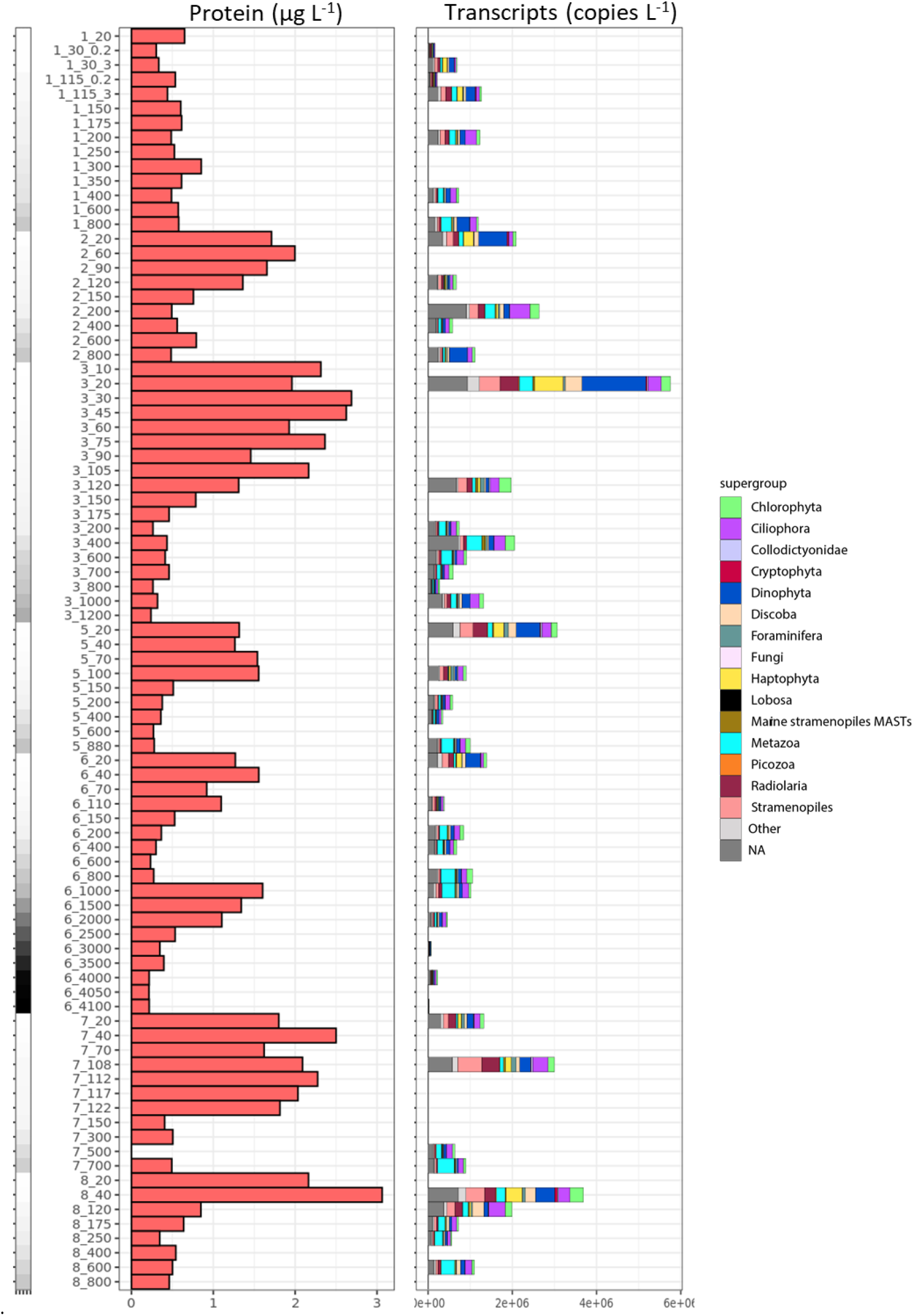
Total extracted protein concentrations (μg L^−1^) and transcript concentrations (copies L^−1^) across stations and depths. Extracted proteins include prokaryotes and eukaryotes (Fig. 2). Transcript concentrations are only shown for the eukaryotic community, since poly-A (eukaryotic) selection of transcripts was performed during RNA processing. For transcripts, the community breakdown of eukaryotic supergroups is shown.

**Supplemental Fig. 17.**
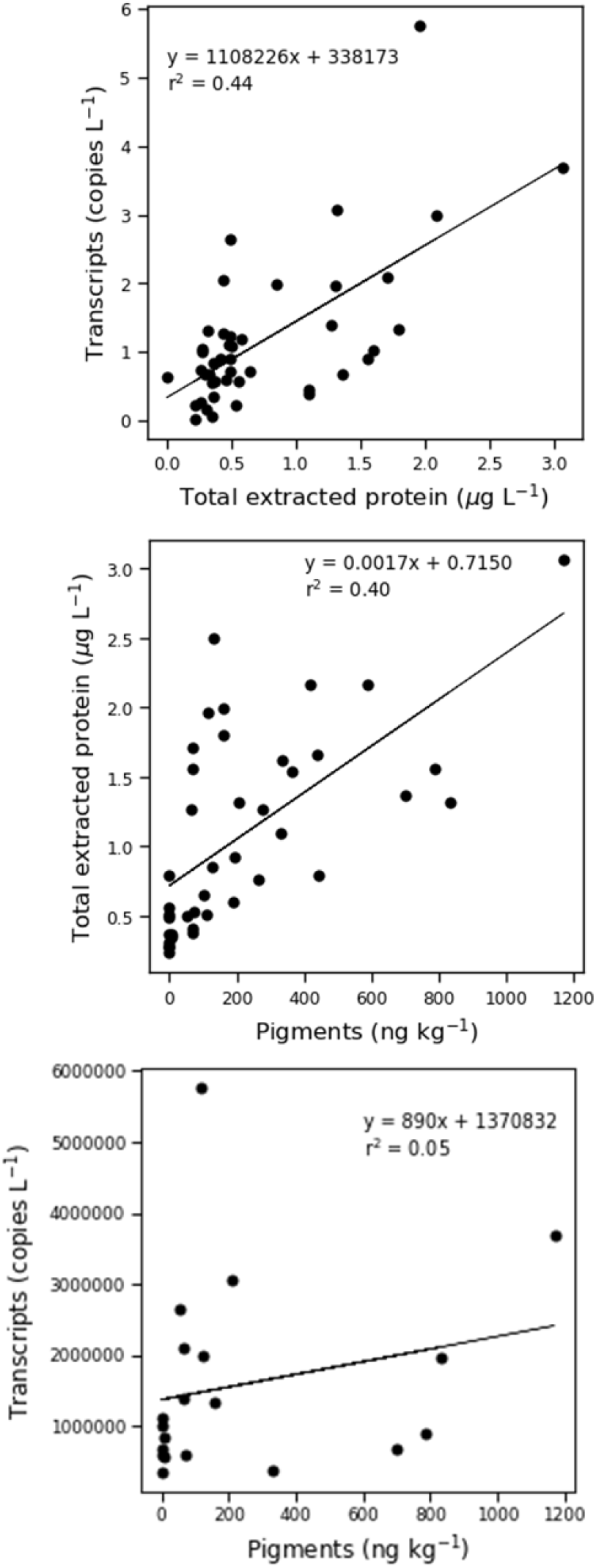
Linear relationships between eukaryotic transcript estimates (copies L^−1^), total extracted protein (μg L^−1^), and total pigment concentrations (ng kg^−1^).

**Supplemental Fig. 18.**
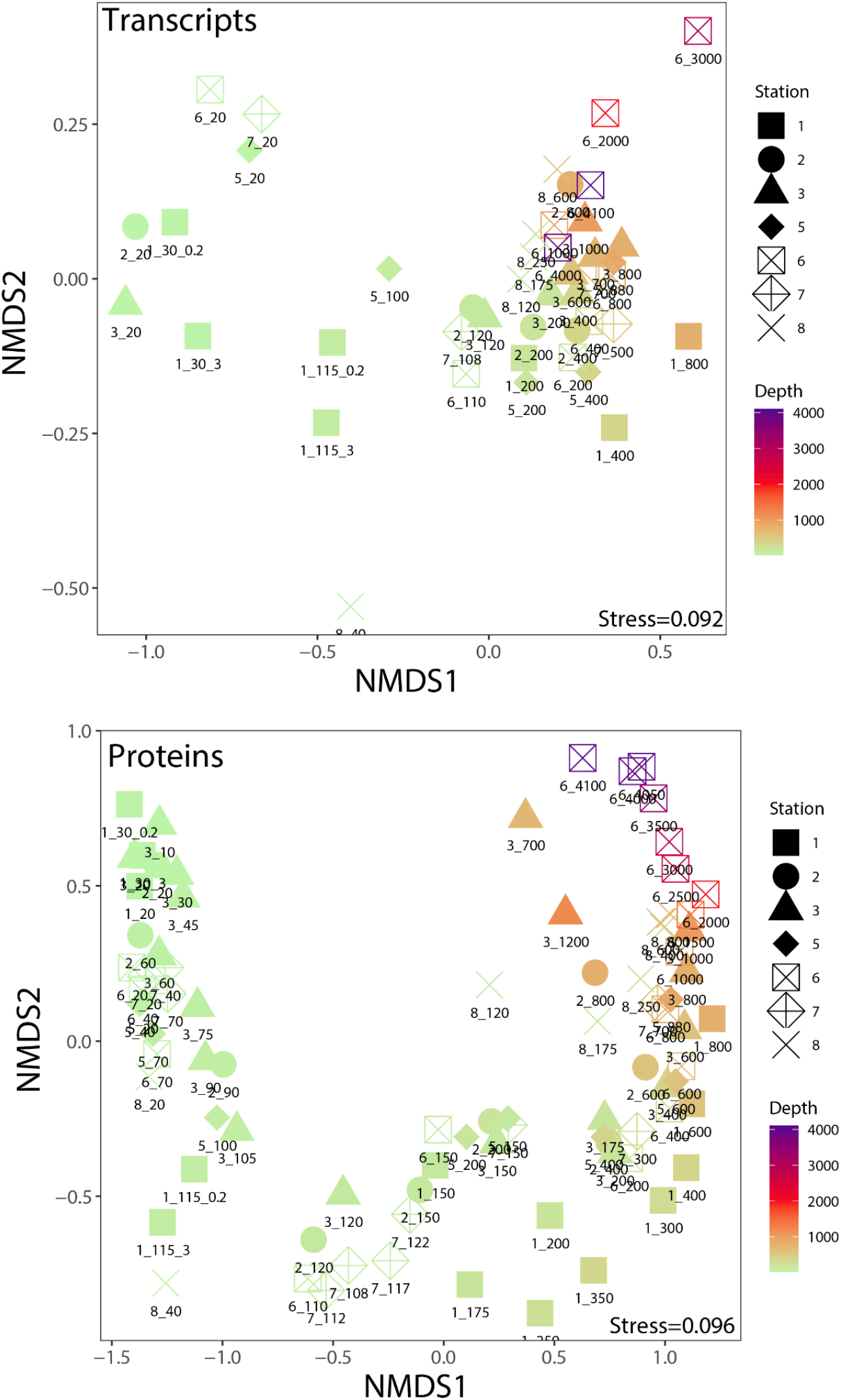
Non-metric multidimensional scaling (NMDS) ordinations using Bray-Curtis distance with community-wide TPM (transcripts) or NSAF (protein) and log-normalized counts. Only ORFs having both a taxonomic and functional annotation were included.

**Supplemental Fig. 19.**
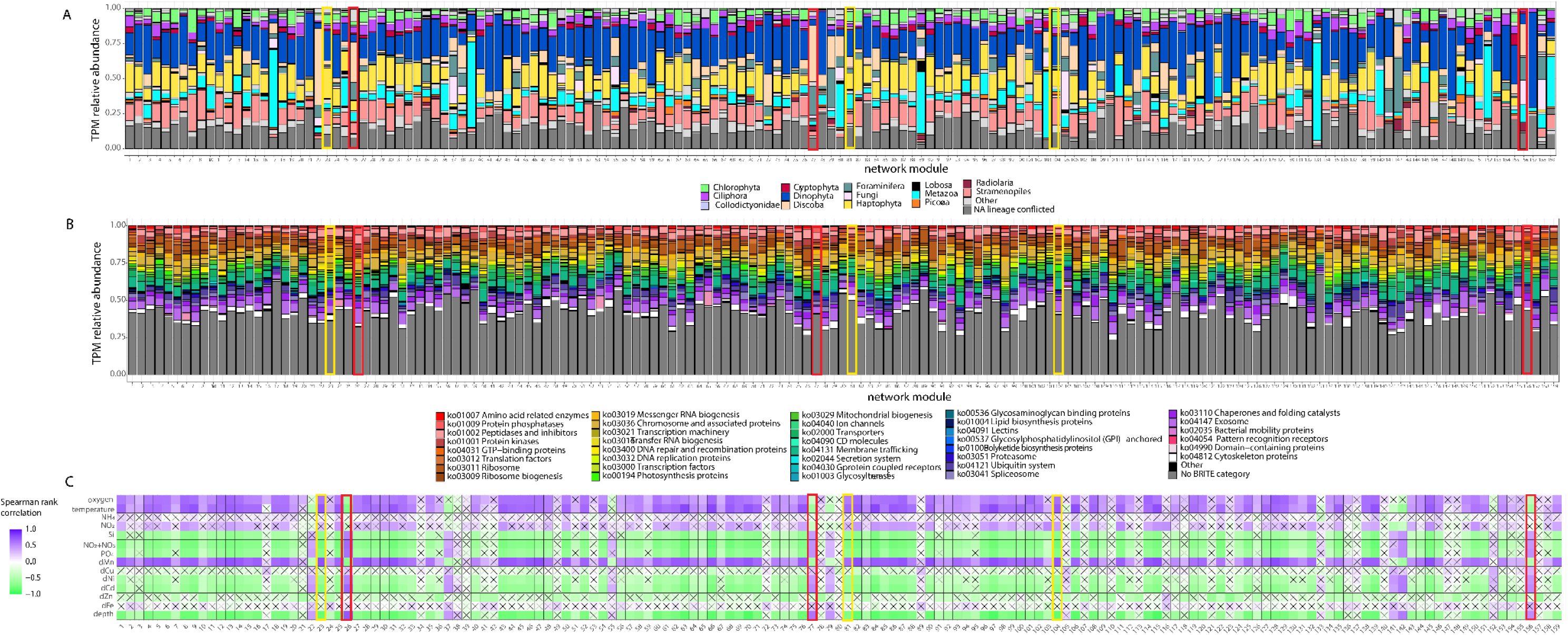
Transcript network built using ORFs with a taxonomic and functional annotation. TPM counts were summed to the supergroup and KEGG annotation level (e.g., Chlorophyta_ko:K00053). Modules were investigated through taxonomic (A) and functional (B) composition. KEGG IDs were grouped into BRITE categories for a broader functional classification. Environmental context for the modules was determined by calculating eigengene values and correlating with environmental data (C). Insignificant Spearman rank correlations are denoted with an “x” (*p* > 0.05).

## Notes

### Competing Interest Statement

The authors have declared no competing interest.

